# It takes tau to tango: Investigating the fuzzy interaction between the R2-repeat domain and tubulin C-terminal tails

**DOI:** 10.1101/2023.02.09.527845

**Authors:** Jules Marien, Chantal Prévost, Sophie Sacquin-Mora

## Abstract

The microtubule-associated protein (MAP) tau plays a key role in the regulation of microtubule assembly and spatial organisation. Tau hyperphosphorylation affects its binding on the tubulin surface and has been shown to be involved in several pathologies such as Alzheimer disease. As the tau binding site on the microtubule lays close to the disordered and highly flexible tubulin C-terminal tails (CTTs), these are likely to impact the tau-tubulin interaction. Since the disordered tubulin CTTs are missing from the available experimental structures, we used homology modeling to build two complete models of tubulin heterotrimers with different isotypes for the β-tubulin subunit (βI/αI/ βI and βIII/αI/βIII). We then performed long timescale classical Molecular Dynamics simulations for the tau-R2/tubulin assembly (in systems with and without CTTs) and analyzed the resulting trajectories to obtain a detailed view of the protein interface in the complex and the impact of the CTTs on the stability of this assembly. Additional analyses of the CTTs mobility in the presence, or in the absence, of tau also highlight how tau might modulate the CTTs activity as hooks that are involved in the recruitment of several MAPs. In particular, we observe a *wrapping* mechanism, where the β-tubulin CTTs form a loop over tau-R2, thus stabilizing its interaction with the tubulin surface and simultaneously reducing the CTTs availability for interactions with other MAPs.

## 1. Introduction

α and β-tubulins are the building blocks of microtubules (MTs), a central element of the cell cytoskeleton, which plays an essential part in cell division, signaling and intracellular transport.^1^ The assembly of α/β-tubulin heterodimers leads to the formation of dynamic tubular fibers. Each tubulin protein is formed of a well-defined globular domain (core), ending with a disordered and negatively charged C-terminal tail (CTT). All cells contain multiple variants (known as isotypes) of α and β-tubulin. While the sequence of the tubulin body is highly conserved, most of the sequence variation between isotypes is located on the CTTs, and it appears that these isotypes are not functionally interchangeable,^2, 3^ as they might modulate the MT assembly,^4, 5^ dynamics^6^ and interactions with other proteins.^7^ In addition, tubulin CTTs are targets for many post-translational modifications (PTMs), and the diversity induced by these genetic and chemical variations is often referred to as the *tubulin code*, and acts as a fine regulator of microtubule function.^8, 9^ In particular, the tubulin tails display different mobility patterns,^10^ which can contribute to the modulation of MT dynamics^11^ and interactions with motor proteins such as kinesin and dynein^12^ and several intrinsically disordered proteins, including tau.^13-16^

Tau is a long (441 amino acids) disordered Microtubule Associated Protein (MAP), which is abundant in brain and neuronal tissues (where it constitutes more than 80% of MAPs).^17^ It regulates the MTs assembly and spatial organization,^18^ and controls the motility of motor proteins on MTs.^19-22^ Tau is involved in numerous neurodegenerative diseases, named tauopathies, including Alzheimer’s disease (AD), which are characterized by fibrillar aggregation.^23^ The binding of tau on MTs, and its propensity to aggregate are affected by PTMs, in particular the hyperphosphorylation of tau has been shown to be involved in several pathologies.^24^ For example, in AD brains, tau is found to be phosphorylated simultaneously on more than 20 positions, mostly serine and threonine residues, and these PTMs impair the protein interaction and its stabilization of microtubules.^25, 26 27^ Due to the disordered nature of tau, the tau/MT assembly is a *fuzzy complex*,^28, 29^ which remains difficult to characterize with classical structural biology approaches. In particular, the disordered CTTs do not appear in the tubulin structures obtained by crystallography or cryo-EM that are currently deposited in the Protein Data Bank^30^ (PDB).

In this perspective, we combined homology modeling and molecular dynamics (MD) simulations to build a complete model, including the CTTs, of the complex formed by the β/α/β tubulin heterotrimer and the tau R2 domain for two tubulin isotypes (βI/αI/βI and βIII/αI/βIII). We used the resulting trajectories to investigate the impact of the tubulin CTTs on the tau/tubulin interaction, and monitor the contacts formed by the partners along time to identify residues that play a key part in the protein interface. In particular, specific attention was payed to tau serine residues, as they are likely to be phosphorylated in tauopathies. We also investigated the CTTs mobility in the presence, or in the absence, of tau-R2 on the tubulin surface in order to understand how tau might modulate the CTTs activity as hooks that are involved in the recruitment of several MAPs.

## 2. Material and Methods

### Building the tubulin C-terminal tails and selecting starting models for the simulations

We used the structure of the complex formed by two β- and one α-tubulin subunits bound with the R2 domain of tau (a 27-residues fragment ranging from Lys274 to Val300) obtained by cryo-EM (pdb: 6CVN)^31^ as a template. As this structure does not account for the tubulin CTTs, we used Modeller v10.2^32, 33^ to build two different systems including different tubulin isotypes, namely βI/αI/βI and βIII/αI/βIII. βI tubulin is the most frequently expressed isotype in all cell types, while βIII tubulin is mainly expressed in neuronal cells and is overexpressed in cancer cells.^34-36^ We used sequences from sheep tubulin (which were chosen to fit experiments from experimental collaborators) for the αI (gene TUBAIA, Uniprot^37^ accession number D0VWZ0) and βI (gene TUBB, Uniprot accession number W5PPT6) subunits, and from human tubulin (since the sheep sequence is not available) for the βIII subunit (gene TUBB3, Uniprot accession number Q13509), see Table 1. The Modeller process is based on a classic comparative modeling method consisting of four sequential steps: template selection, template-target alignment, model building and a final model evaluation. We used the Clustal^38^ tool for sequence alignment to determine the homology percentage between the target sequences and the 6CVN-template sequence. Because of the high similarity between the target sequences and the structural template (over 94% sequence identity for the tubulin core fragments, see Table SI-1), there was no need for a manual rearrangement of the alignments.

**Table 1:** Aligned sequences of the tubulin C-terminal tails.

| Isotype | Sequence | Length |
| --- | --- | --- |
| αI (D0vWZ0) | SVEGEGEEEGEEY | 13 |
| βI (W5PPT6) | DATAEEEE--DFGEEAEEEA | 18 |
| βIII (Q13509) | DATAEEEG--EMYEDDEEESEAQGPK | 24 |

To select non-redundant CTT structures as starting points for the simulations, we first generated 100 models with Modeller, where only the CTTs structures would vary. In a second step, we used the gmx_cluster tool from the GROMACS^39^ suite, with clustering cutoffs on the CTT backbones of 8.5 Å and 9.5 Å, for the βI/αI/βI and βIII/αI/βIII systems respectively. For both systems, we selected a representative structure from the two most populated clusters. In addition, an outlier structure (identified by increasing the cutoffs to 10 Å and 11 Å respectively) was also selected for both systems. These sets of three structures for each one of the two systems were used as seeds for the MD simulations to efficiently sample the CTTs conformational space and are shown in Figure 1. In addition, we removed R2 from these six models to create seed structures for tubulin heterotrimers with no tau-R2 bound on the tubulin surface. We also cleaved the tubulin CTTs in the three models built for the βI/αI/βI assembly (namely models 17, 22 and 84) to investigate the interaction of tau-R2 with a tubulin heterotrimer without CTTs. Finally, the tau-R2 structures from the βI/αI/βI models were isolated in order to study the behavior of tau-R2 in solution. Table 2 provides a summary of all the systems built for the present study.

**Figure 1:**
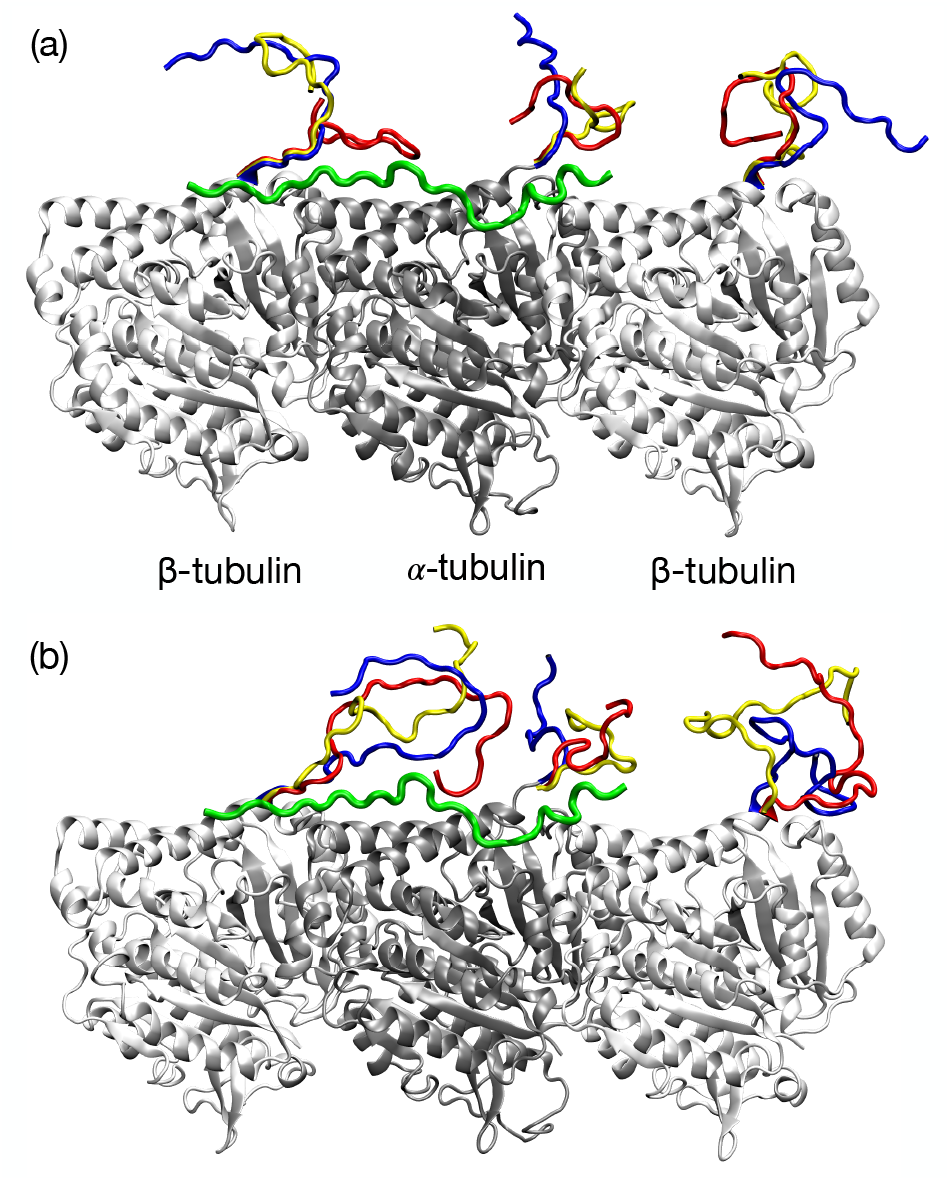
Starting structures for the tau-R2/tubulin assembly. The tubulin folded core is shown in dark and light grey for the α and β subunits respectively, while the disordered CTTs are shown in red, blue and yellow, and tau-R2 is shown in green. (a) βI/αI/βI isotype (b) βIII/αI/βIII isotype. Figures 1, 2, 5 and 6gh were prepared using Visual Molecular Dynamics.^45^

**Table 2:** Summary of the MD simulations performed in the study. For each setup (column), we ran three replicas of 200 ns.

|  |  | Box size | Protein atoms | Ions | Water molecules |
| --- | --- | --- | --- | --- | --- |
| tau-R2 in solution |  | 100x100x100Å<sup>3</sup> | 446 | Na<sup>+</sup> : 89<br>Cl<sup>-</sup> : 93 | 31127 |
| Tubulin heterotrimer without tau-R2 | βI/αI/βI isotype | 130x130x190Å<sup>3</sup> | 20585 | Na<sup>+</sup> : 340<br>Cl<sup>-</sup> : 266 | 93380 |
|  | βIII/αI/βIII isotype | 140x140x215Å<sup>3</sup> | 20781 | Na<sup>+</sup> : 431<br>Cl<sup>-</sup> : 357 | 125429 |
| tau-R2/tubulin complexes | Tubulins without CTTs | 130x130x190Å<sup>3</sup> | 20377 | Na<sup>+</sup> : 308<br>Cl<sup>-</sup> : 267 | 93635 |
|  | βI/αI/βI isotype | 130x130x190Å<sup>3</sup> | 21031 | Na<sup>+</sup> : 336<br>Cl<sup>-</sup> : 266 | 92318 |
|  | βIII/αI/βIII isotype | 140x140x215Å<sup>3</sup> | 21227 | Na<sup>+</sup> : 426<br>Cl<sup>-</sup> : 356 | 125253 |

### All-atom Molecular Dynamics simulations

#### Simulation setup

All systems (see Table 2) were prepared using the solution builder module from the CHARMM-GUI^40^ (https://www.charmm-gui.org/) input generator. The complete system was solvated with TIP3P water molecules^41^ in rectangular boxes (with a larger box for the βIII/αI/βIII system accounting for the longer CTT of tubulin-βIII), in a physiological solution of NaCl (0.15 mol.L^-1^), with amino-acids protonation states set to pH 7 (and unprotonated histidine residues). We used NAMD v2.14,^42^ with the CHARMM36m^43^ force-field under periodic boundary conditions with the Particle Mesh Ewald (PME) method,^44^ and covalent bonds involving hydrogen atoms were constrained using the SHAKE algorithm.^45^ We used a time step of 2 fs, and data acquisition was set every 100 ps. The simulation procedure starts with 10000 steps of energy minimization and 2 ns of equilibration in the NVT ensemble, which are followed by 200 ns of production in the NPT ensemble at 300K. The temperature was simulated through Langevin dynamics, and pressure was set at 1 atm using a Nose-Hoover Langevin Piston algorithm.^46, 47^ The nonbonded interaction cutoff was set to 1.2 nm, and the Particle Mesh Ewald algorithm^48^ was used for electrostatic calculations with a grid space of 1 Å. We used the ColVar module^49, 50^ to constrain the α-carbons of the tubulin cores by a harmonic bias, with a scaled normalized force constant of 0.05 kcal.mol^-1^ based on a collective variable on the overall orientation angle of the complex, in order to avoid a rotation of the system that would lead to self-interaction with the image systems from the periodic boundary conditions. This protocol was repeated for each of the three initial structures generated for the R2/tubulin complex (without CTTs, with the βI/αI/βI CTTs, and with the βIII/αI/βIII CTTs). Altogether we have 3*200 ns of production in the NPT ensemble for each complex, and for the systems with the tubulin heterotrimer alone (without tau-R2).

In addition, we also performed a MD simulation of tau-R2 in water (in a cubic water box of 10 nm side length) following the same protocol and using the three initial tau-R2 structures taken from the complex with the βI/αI/βI isotype.

#### Analysis

For the systems that include a tubulin heterotrimer, all trajectories were first aligned on the tubulin core in post-processing. We used VMD Analysis^51^ tools and ad-hoc Python scripts using the MDAnalysis^52, 53^ and Mdtraj^54^ libraries to analyze the MD trajectories. MMPBSA binding enthalpies between R2 and the tubulin heterotrimer, taking into account the effect of the solvent, were calculated by combining the last 150 ns, with a 10-frame stride, of each one of the three models. To perform the calculations, we used Cafe_plugin,^55^ a VMD plugin, and the APBS webserver.^56^

## 3. Results and discussion

### Conformational variability of the disordered fragments

A common issue with classical force fields used for the modeling of proteins is that they tend to overstabilize secondary structure elements,^29^ thus biasing the observation of the unfolded states that characterize disordered protein fragments.^57, 58^ We chose to use the Charmm36m force-field for our MD simulations, as it was specifically developed to address this problem. In addition we calculated DSSP^59^ (for Definition of Secondary Structure of Protein) graphs, which highlight the most likely secondary structure adopted by each residue given the global 3D structure of a protein, for the tau-R2 fragment (Figure SI-1) and the tubulin CTTs (Figure SI-2).

The DSSP graphs resulting from the simulation of tau-R2 in solution highlight the disordered nature of the protein, as they display only transient secondary structure elements, which vary from one trajectory to the other (see Figure SI-1a-c). This lack of stability for the tau-R2 structure in solution is also visible in the RMSD plots shown in Figure SI-3. The DSSP graphs for the tubulin-bound R2 show no secondary structures for the N-ter part of tau-R2. The last five residues in the C-ter part (Asp295-Val300) can form a helical structure but only transiently (for less than 15ns, see Figure SI-1d-l), which can be related to the formation of helical structures observed by Li et al.^60^ in tau repeat regions when binding to tubulin. During one of the simulations of tau-R2 bound on tubulins without CTTs (model 22), the C-ter fragment of tau will detach from the tubulin surface and fold upon itself, thus forming a β-bridge between Val287 and Val300 (see Figure SI-1e, the RMSD profile of tau-R2 in Figure SI-4g, and the movie of the trajectory provided in the Supplementatry information).

### Investigating the tau-R2/tubulin interface

#### Electrostatic properties of tau-R2 and the tubulin heterotrimer

The surface electrostatic potentials for tau-R2 and both isotypes of the tubulin heterotrimer were calculated separately using the structures in the complex and are shown on Figure 2. While tau-R2’s surface is mostly positively charged due to six lysine residues (see inserts in Figure 2), the tubuline’s surface displays a broad electronegative patch all around the tau-R2 binding site (see the upper panels of Figures 2a and b). In addition, the CTTs are also negatively charged, as they contain many glutamate residues.

**Figure 2:**
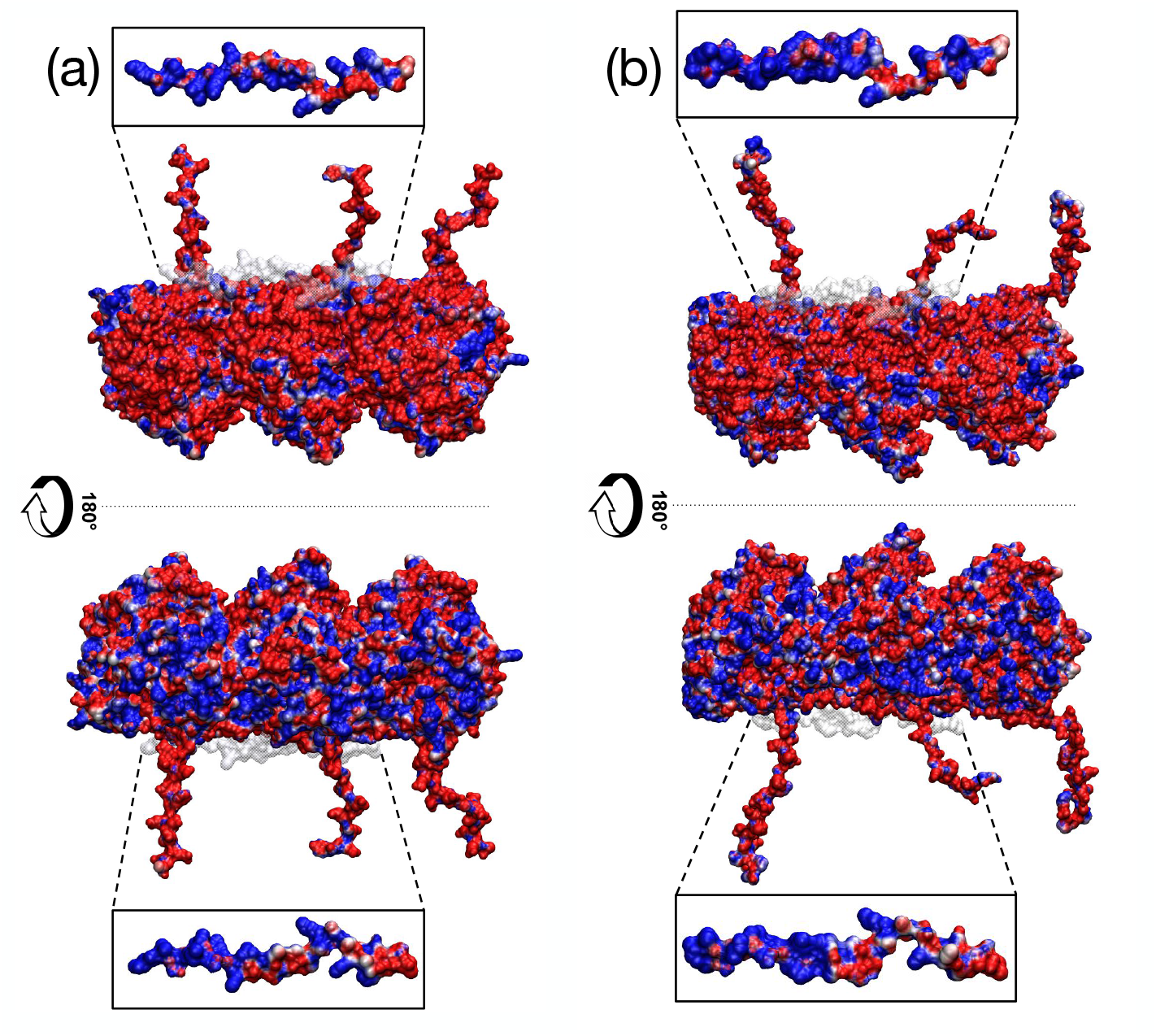
Surface electrostatic potential for the tubulin heterotrimer and tau-R2 (shown in the inserts), the complex orientation in the upper panels is the same as in Figure 1. Positive potentials are colored in blue and negative potentials in red. (a) βI/ αI/βI isotype (frame 521 from model 17 trajectory) (b) βIII/αI/βIII isotype (frame 1142 from model 16 trajectory). These two frames were selected for clarity as the CTTs extended conformation allows a better visualization of the surface electrostatic properties.

#### Conformational stability of the tau-R2/tubulin assembly

The complex overall structure remains stable during all the trajectories, both without and with CTTs, as shown by the RMSD profiles in Figure SI-4. However, this stability is mostly due to the folded tubulin cores (see panels d-f in Figure SI-4), as both the CTTs and the tau-R2 fragment display much higher RMSDs during the trajectory (panels g-k in Figure SI-4), thus highlighting their flexible and disordered nature.

Figures SI-5a-c show the evolution of Fnat, the fraction of native contacts (i.e. residues from the initial tau-R2/tubulin interface with heavy atoms less than 5 Å away) as a function of time for the tau-R2/tubulin interface. Fnat remains in the [0.4-0.6] range for all trajectory except for one replica (model 84 from the βIII/αI/βIII isotype), a value lower than those observed for protein complexes involving fully folded partners,^61^ thus characterizing the fuzzy nature of the tau/tubulin interface.^29^ Despite the decrease of the Fnat value, the total number of contacts formed between tau-R2 and tubulin remains stable along time (as shown in Figures SI-5d-e), with a slightly larger average value in the case of the systems including the tubulin CTTs, as these provide opportunities for more interactions with tau-R2 during the simulations.

Meanwhile, Figure 3a shows the average RMSF values for the tubulin-bound tau-R2 fragment. The profiles highlight a stable binding core centered on residue Ser289, which corresponds to the R2 strong interacting region identified by Brotzakis et al.^62^ in their study integrating Cryo-EM data into MD simulations. Interestingly, the RMSF values can be larger in the case of the systems including tubulin CTTs (see Figures 3 and SI-6) in particular for the N- and C-ter ends of tau-R2, which can interact with the CTTs from the first (starting from the left of Figure 1) β-tubulin and the α-tubulin subunits. These interactions then lead to a lift of tau-R2 from the tubulin-core surface and an increased RMSF, either on the N-terminal side of tau-R2 (see the trajectories starting from models 17 and 38 in Figures SI-6b-c), or on its C-terminal side (trajectories starting from models 22 and 16 in Figures SI-6b-c)

**Figure 3:**
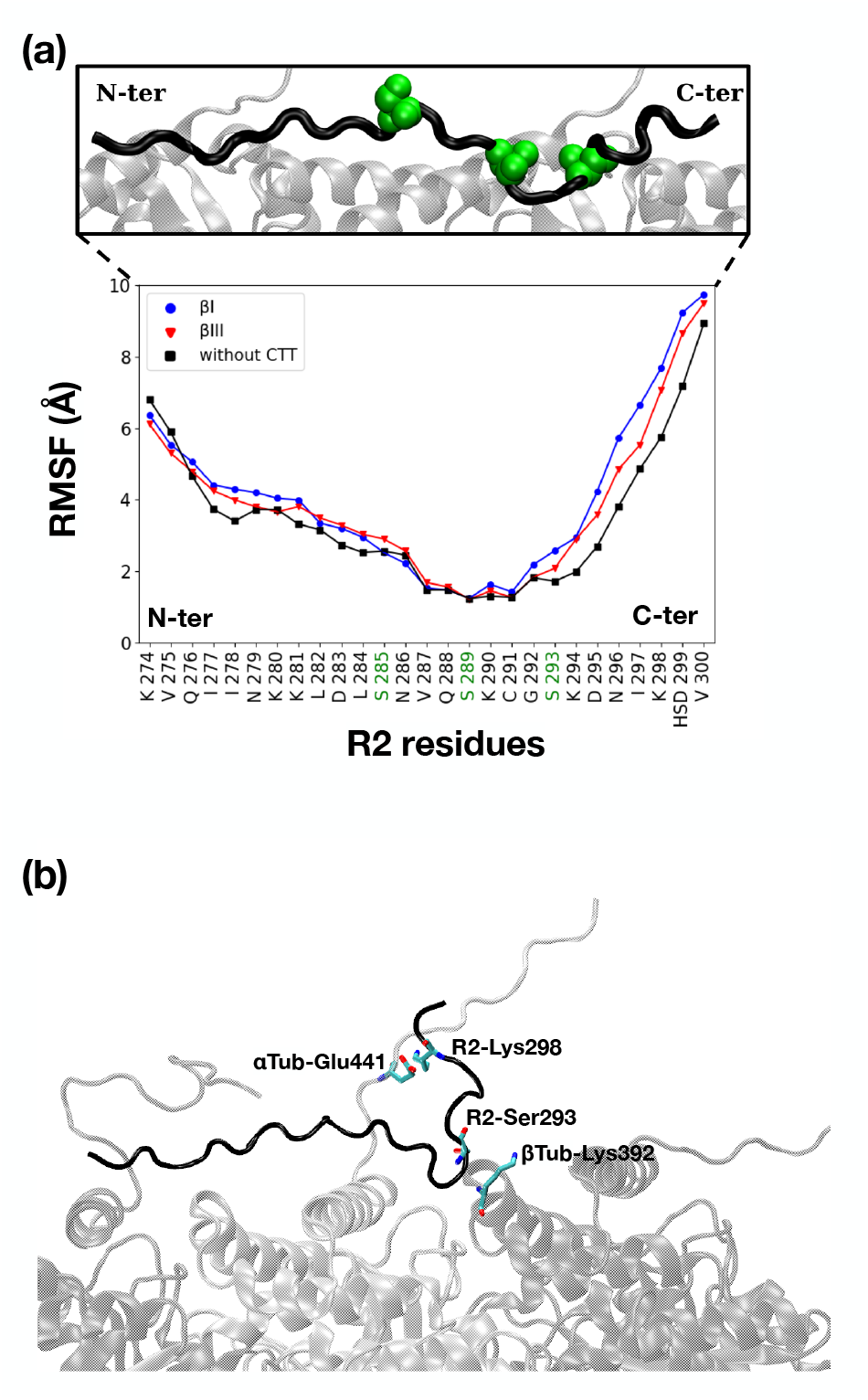
(a) RMSF of tau-R2, average value over the three simulations for each system. The three serine residues that are phosphorylated in tauopathies are highlighted in green along the sequence and shown as van der Waals spheres in the upper panel. (b) Snapshot from the trajectory starting with model 22 (for the βI/αI/βI isotype), where the R2-CTT (in black) is lifted from the tubulin surface.

Finally, the binding enthalpies between tau-R2 and the tubulin are available in Table 3.

**Table 3:** Average binding enthalpies (in kcal.mol^-1^), with the different contributions, between tau-R2 and the tubulin heterotrimer.

| tau-R2/tubulin complex | Total binding enthalpies | $\Delta E_{el}$<br>(electrostatic) | $\Delta E_{vdw}$<br>(van der Waals) | $\Delta G_{p}^{sol}$<br>(PB term) | $\Delta G_{np}^{sol}$<br>(SASA term) |
| --- | --- | --- | --- | --- | --- |
| Tubulins without CTTs | -59.5 ± 18.6 | -2236.4 ± 199.4 | -73.3 ± 12.0 | 2261.3 ± 196.7 | -11.1 ± 1.5 |
| βI-αI-βI complexes | -74.9 ± 22.0 | -3132.1 ± 258.7 | -79.0 ± 18.9 | 3148.6 ± 250.3 | -12.4 ± 2.7 |
| βIII-αI-βIII complexes | -74.1 ± 20.1 | -3197.5 ± 232.1 | -73.7 ± 15.4 | 3209.0 ± 234.6 | -11.9 ± 2.2 |

The total values are mostly due to the electrostatic term of the potential (with a van der Waals contribution accounting for less than 5% of the electrostatic term), and concur with an earlier computational work on a similar system (unphosphorylated tau-R2 bound to a tubulin heterotrimer without CTTs) using a different force field (ff14SB) by Man et al.,^63^ thus showing the robustness of MD simulations for the investigation of protein assemblies. On Table 3, we can see how the tubulin CTTs help stabilizing the system by increasing the global binding energy by roughly 25%, in agreement with earlier experimental data obtained by FRET^64^ and fluorescence correlation spectroscopy.^65^

#### Detailed view of the contacts between tau-R2 and the tubulin heterotrimer

Figure 4 shows a map of the contacts formed between tau-R2 and the tubulin heterotrimer during the simulations, the three-dimensional distribution of the contact points over the tubulin surface is shown in Figure 5. The contacts between tau-R2 and the tubulin cores are highly conserved in the three systems, and for both tubulin isotypes, the residues from the CTTs of first β-subunit (on the left of Figure 5b-c) and α-subunit are also involved in the interaction with tau-R2. In the R2 strong interacting region, residues from the α- subunit forming long lasting contacts, such as Glu423, Ala426, Ala427 and Lys430 correspond to tau binding residues identified using NMR spectroscopy by Lefevre et al.^66^These tubulin residues also concur with the binding pattern identified by Jimenez^67^ using Gaussian-accelerated MD simulations with a different force field (Amber ff14SB^68^) on a tubulin-tau assembly that did not include tubulin CTTs. Furthermore, tubulin forms very stable contacts with tau-R2 around its three serine residues, Ser285, Ser289 and Ser293, which have been identified as phosphorylation sites in tauopathies.^24-27^ We also investigated hydrogen bonds formed by the serine residues within tau-R2 and with the tubulin heterotrimer (see Figure SI-7). Ser289 forms a very stable hydrogen bond with Glu434 from the tubulin α-subunit. This interaction is further reinforced when adding the CTTs, as the tubulin αCTT interacts with the positively charged C-ter of tau-R2 and in particular prevents residue Lys298 to interact with αTub-Glu434 (see Figure 4). Inversely, adding the tubulin CTTs destabilizes the transient hydrogen bond formed between tau-R2 Ser293 and Lys392 from the second β-subunit (when reading Figure 5 from the left), since the C-ter of tau-R2 will now be lifted from the tubulin surface to interact with the tubulin αCTT (see the snapshot in Figure 3b). This phenomenon also leads to the RMSF increase shown in Figure SI-6.

**Figure 4:**
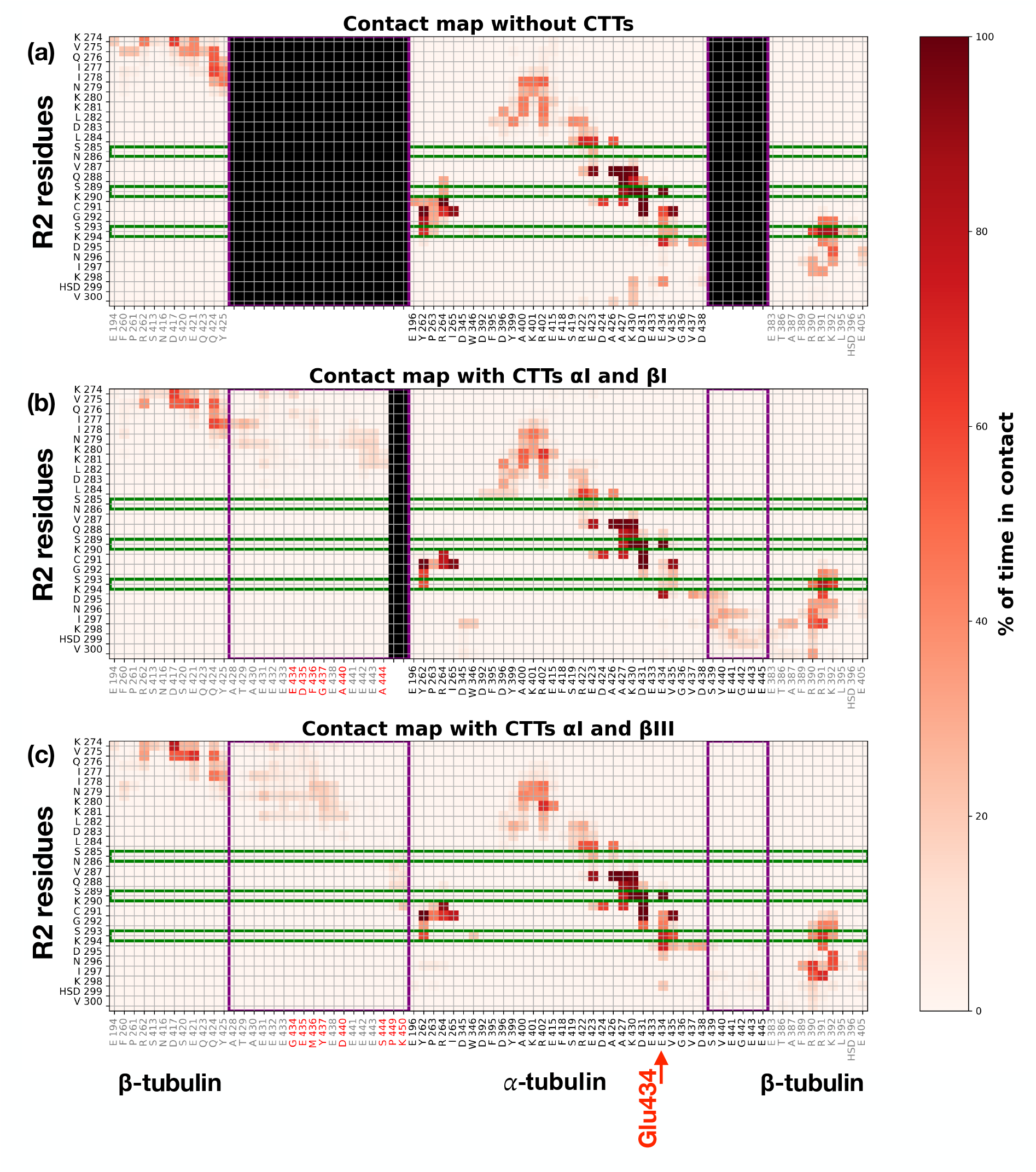
Contact maps for the tau-R2/tubulin interface. The map only lists tubulin residues that are in contact with tau-R2 during at least 10% of the cumulated 600 ns from the three MD trajectories ran for each system. The tubulin CTTs for the βI/αI/βI and βIII/αI/βIII isotypes (central and lower panels) are delimited by a purple frame, while the potentially phosphorylated serines (Ser 285, 289 and 293) of tau-R2 are highlighted by green frames. (a) Tubulin without CTTs (b) βI/αI/βI isotype (c) βIII/αI/ βIII isotype.

**Figure 5:**
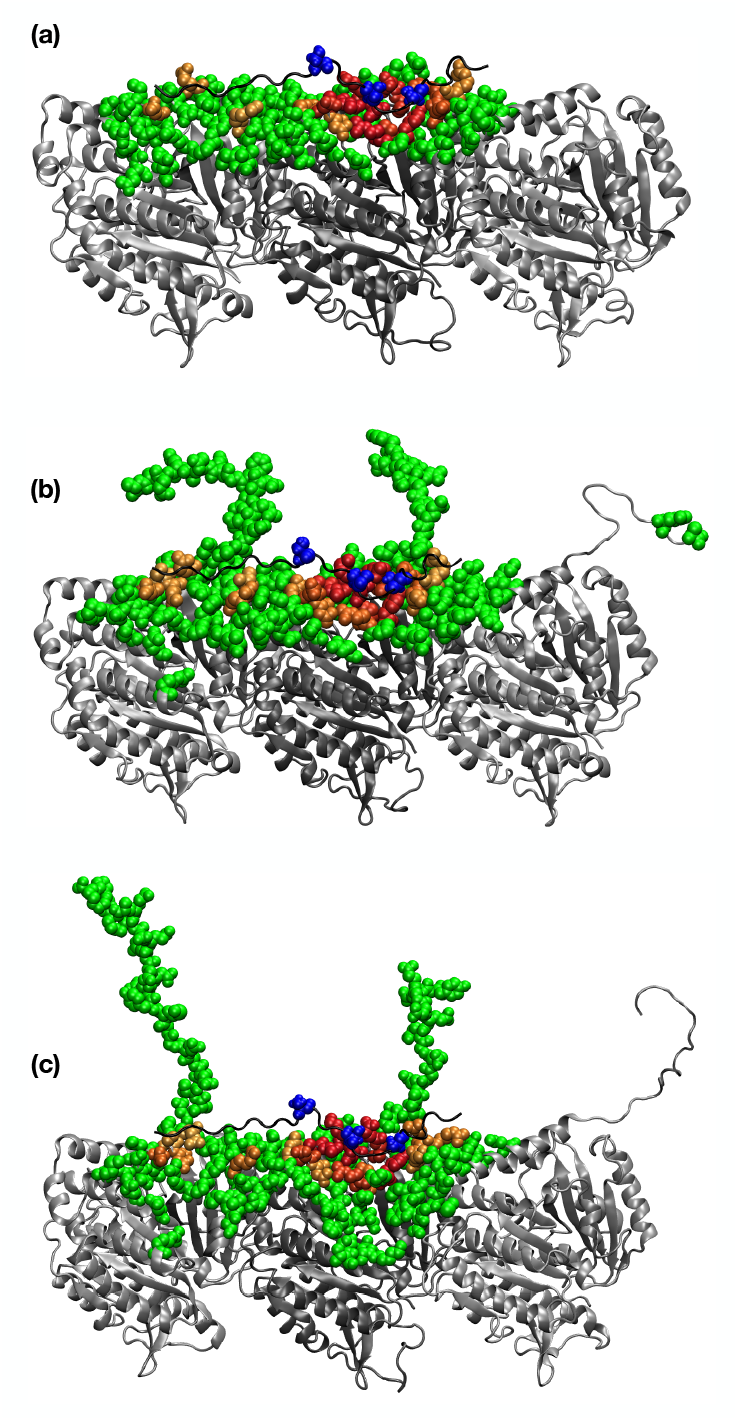
3-D distribution of the tubulin residues forming contacts with tau-R2 during the MD simulations. Transient contacts (less than 50% of the trajectory) are shown as green van der Waals spheres, while long lasting contacts are highlighted with a color scale ranging from orange (contacts formed during 50% of the trajectory) to red (permanent contacts). Serines 285, 289 and 293 of tau-R2 are shown as blue van der Waals spheres. (a) Tubulin without CTTs (b) βI/αI/βI isotype (c) βIII/αI/βIII isotype.

In addition to the R2 strong interacting region, Figures 4 and 5 also highlight a secondary tau binding site on the tubulin surface, located near the interface between the first β-tubulin and the α-tubulin subunits, and which concurs with the weakly interacting region R2w centered around R2-Lys280 and R2-Lys281 defined by Brotzakis et al.^62^ Tau-R2 residues 274-281 from this R2w region were also shown to play an important part in microtubule binding by NMR spectroscopy studies,^69^ and in microtubule dynamics in an experimental study by Panda et al.^70^

#### Impact of tau-R2 on the CTTs mobility pattern

Tubulin CTTs play an important part in the regulation of MT stability, in particular through their mobility and conformational dynamics.^71, 72^ As they protrude from the MT surface, they are sometimes labeled as *E-hooks*,^73^ due to their rich content in negatively charged glutamic acid residues, and are involved in the recruitment of several MAPs.^74-76^ Since the tau binding site on the tubulin surface lies close to the CTTs, the interaction between tau and tubulin is likely to have an effect on the CTTs mobility. To investigate this issue, we also ran MD simulations (following the protocol described in the Material & and Methods section) for the βI/αI/βI and βIII/αI/βIII isotypes of the tubulin heterotrimer, but this time without tau-R2. The CTT mobility patterns for both isotypes during the trajectories without and with tau-R2 are shown in Figure SI-8 via the distribution of the position of the CTTs center of mass (COM), using the same representation mode as in ref. 10. Interestingly, the CTT of the first β-subunit displays the same shift toward the surface of the α-subunit that was already observed in ref. 10. In addition the αI-CTT mostly interacts with the tubulin surface close to the tubulin longitudinal polymerization interface (between the α-subunit and the second β-subunit), in agreement with an earlier study of Chen et al.,^72^ which used both experimental and simulation approaches. Comparing the mobility patterns without and with R2, we can see how the binding of tau-R2 on the tubulin surface seems to prevent the interaction between the tails of the α- subunit and the second β-subunit, with the αI-CTT slightly shifting its COM distribution toward the surface of the α-subunit.

For both tubulin isotypes, we also monitored the number of contacts formed between each one of the three CTTs and the tubulin core, or between the CTTs and the tubulin core and tau-R2, during the trajectories without and with bound tau-R2 on the tubulin surface respectively. The resulting density plots are shown in Figure s6a-f and SI-9, while the number of contacts for each CTT as a function of time is available in Figures SI-10 and SI-11. For the simulations without tau-R2, and for both the βI/αI/βI and βIII/αI/βIII isotypes, the CTT of the first β-subunit presents a bimodal distribution (see Figure 6a) with a predominant, low contact number state (termed as the *free state*) and a high contact number state (termed as the *bound state*). In the free state, the CTT/ tubulin core contacts are only formed via residues lying at the base of the CTT, while in the bound state all the CTT residues can form contacts with the tubulin core or with R2 (as shown in the snapshots in Figure 6g and h). Adding tau-R2 on the tubulin surface leads to a clear shift of the distribution in favor of the CTT bound state for both isotypes (see Figure 6d). In the case of the α-CTT (from the central tubulin subunit) the impact of tau binding on the contact number distribution is more important for the βI/αI/βI isotype (blue lines in Figures 6b and e). For the contacts formed by second β-subunit CTT, the βIII/αI/βIII isotypes appears more sensitive to the introduction of tau-R2 in the system (red lines in Figures 6c and f). However, one should keep in mind that only the base of this CTT can form contacts with tau-R2 (see Figures 4 and 5c), as it is located right at the end of the tubulin heterotrimer.

**Figure 6:**
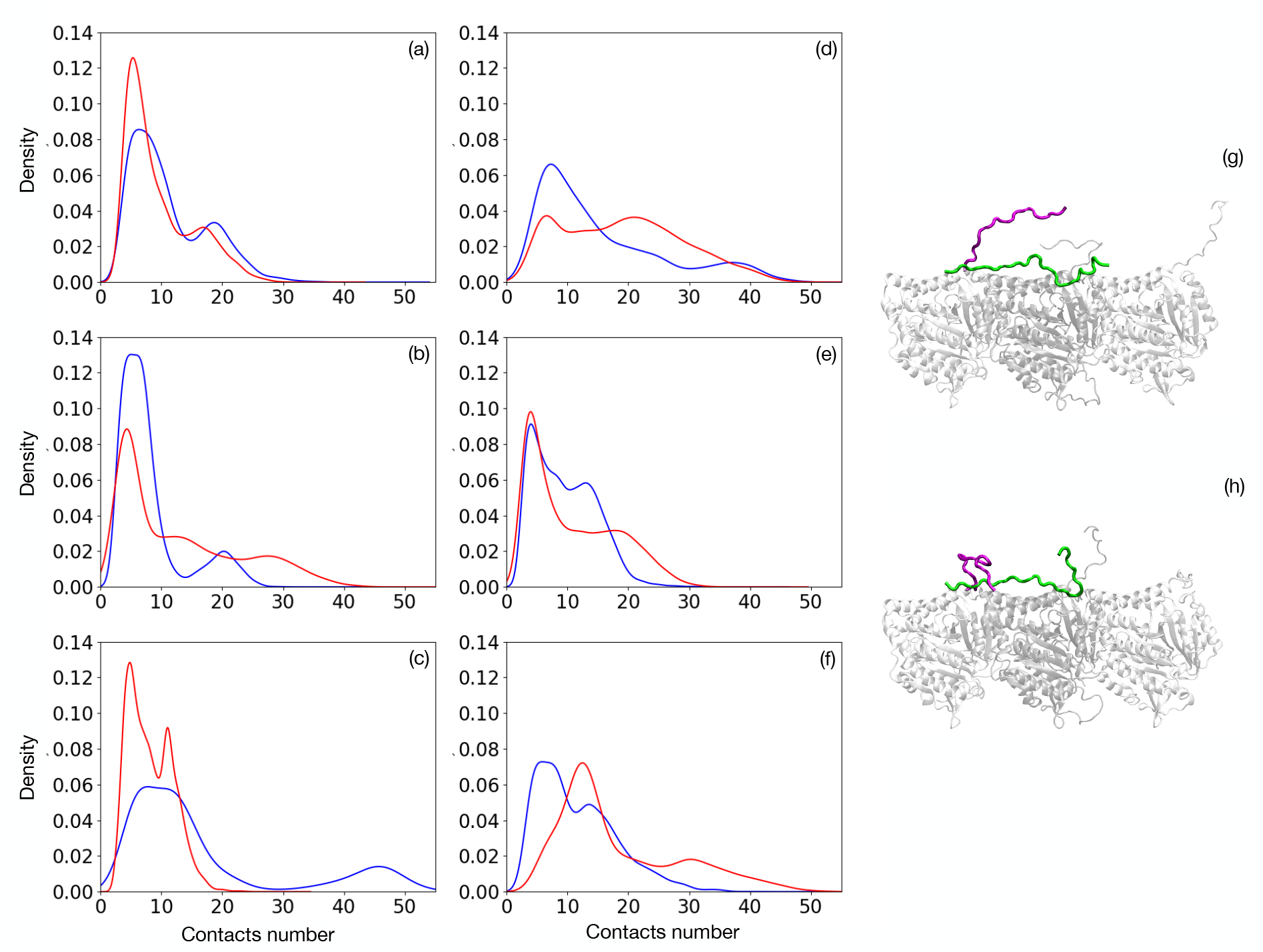
Density for the number of contacts formed by each CTT in the βI/αI/βI isotype (blue lines) and the βIII/αI/βIII isotype (red lines): Simulations without tau-R2, contacts formed between the CTT and the tubulin core (a) First β-subunit, (b) α-subunit (c) second β-subunit.\ Simulations with tau-R2, contacts formed between the CTT and the tubulin core or between the CTT and tau-R2 (d) First β-subunit, (e) α-subunit (f) second β-subunit. Snapshots from the trajectory with the βI/αI/βI isotype and bound tau-R2 starting from model 22. The CTT from the first β-subunit is higlighted in purple, (g) free CTT state,bound CTT state

On several occasions, we could observe for both βI and βIII CTTs a specific type of bound state, where the CTT will wrap around the N-ter extremity of R2 (see Figure 6h), thus stabilizing this weak contacts zone, and resulting in the lowered RMSD of R2 for model 22 from the βI/αI/βI isotype in the 120-200 ns range (see Figure SI-4h) or for model 38 from the βIII/αI/βIII isotype in the 30-200 ns range (see Figure SI-4i). On the residues scale both tubulin isotypes seem to behave differently: the CTT from the first βI-tubulin subunit will form salt bridges with R2-Lys280 and Lys282 via its terminal glutamate residues, which results in the very low RMSF values obtained for the N-ter fragment of R2 (see Figure SI-6). On the other hand the CTT from βIII-tubulin loops over R2 to form a salt-bridge between its terminal Lys450 and residues Glu420 and Glu423 from the α-tubulin subunit. Overall this “wraping mechanism” could in part explain why Tau’s affinity for MTs decreases when CTTs are removed.^14^

## 4. Conclusions

Tau is a disordered protein that plays a key role for the stabilization of microtubules and fine-tunes their interaction with other MAPs. Here, we used a combination of modeling approaches to investigate the binding modes of the tau-R2 repeat on a tubulin heterotrimer on the atomic level. In particular, the reconstruction and explicit modeling of the tubulin C-terminal tails (which are not included in the experimental structures available for the tau/tubulin system) for two different β-tubulin isotypes enable us to see how the tau-R2/tubulin assembly forms a fuzzy complex, with conformational variability arising both from tau-R2 and the disordered CTTs. We also show how these CTTs contribute to stabilizing the assembly despite bringing additional mobility in the system. Our detailed investigation of the tau-R2/tubulin interface highlights the central part played by the three serine residues of tau-R2, and in particular Ser289, which lies in the center of the R2 strongly interacting region of tau. In the future, we plan to explicitly model the phosphorylation of these serine residues and its impact on stability of the tau-R2 interface, in order to better understand on the molecular scale how tau hyperphosphorylation can lead to various pathologies by disrupting its interaction with microtubules.^63, 77^ While numerous studies highlight the role played by tubulin CTTs or tau as regulators of motor proteins motility on microtubules, very few consider both elements (the CTTs and tau) simultaneously. Our results regarding the CTTs mobility and contacts without or with tau-R2 show a strong interplay between the two partners. The shift in favor of the CTTs bound state induced by tau that we observe in our simulations supports the hypothesis formulated by Lessard and Berger^78^ that tau-mediated kinesin regulation might be linked to a competitive interaction between tau and the motor protein for the CTTs. Consequently, further work will also investigate how tau phosphorylation can impact the tubulin CTT’s mobility, since it has been shown experimentally to modulate kinesin motility along microtubules,^21^ a phenomenon where the CTTs are likely to play a part.

## Acknowledgments

This work was supported by the ANR (MAGNETAU-ANR-21-CE29-0024) and by the “Initiative d’Excellence” program from the French State (Grant “DYNAMO”, ANR-11-LABX-0011-01). Simulations were performed using the HPC resources from LBT/HPC.

## Supporting Information Available

Additional data regarding the tubulin modelling (DSSP of the disordered fragments, RMSD on bulk tau-R2, RMSDs of the tau-tubulin assembly, fraction of native contacts and total number of contacts, RMSF of the MT-bound tau-R2, Energies as a function of time, H-bonds formed by the serines in tau-R2, CTT mobility patterns on the MT surface, number of contacts formed by the CTTs) is available as a pdf file. All MD trajectories without solvent and their topologies, and movies for each simulation are deposited in Zenodo (https://zenodo.org/record/7643534#.Y-zy7OzMKzl)

## TOC image

For Table of Contents use only

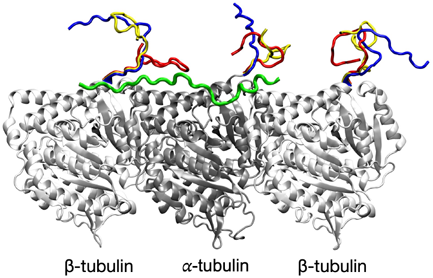

## Notes

https://zenodo.org/record/7643534#.Y-zy7OzMKzl

### Competing Interest Statement

The authors have declared no competing interest.

### Summary of Updates

The article has been clarified, with several figures moved to the supplementary information, and additional MMPBSA calculations for the tau/tubulin binding energy.

