## Supplementary data for "It takes tau to tango: Investigating the fuzzy interaction between the R2-repeat domain and tubulin C-terminal tails"

**Table SI-1:** Homology percentage and sequence alignments between the template sequence (from pdb 6CVN) and the sheep and human tubulins used in the study.

| | αI (TUBAIA) | βI (TUBB) | βIII (TUBB3) |
|---|---|---|---|
| 6CVN (α subunit) | 99.5% | | |
| 6CVN (β subunit) | | 97.9% | 94.4% |

```
alpha1_core_sheep      MRECISIHVGQAGVQIGNACWELYCLEHGIQPDGQMPSDKTIGGGDDSFNTFFSETGAGK
alpha1_6CVN            MRECISIHVGQAGVQIGNACWELYCLEHGIQPDGQMPSDKTIGGGDDSFNTFFSETGAGK
                       ************************************************************

alpha1_core_sheep      HVPRAVFVDLEPTVIDEVRTGTYRQLFHPEQLITGKEDAANNYARGHYTIGKEIIDLVLD
alpha1_6CVN            HVPRAVFVDLEPTVIDEVRTGTYRQLFHPEQLITGKEDAANNYARGHYTIGKEIIDLVLD
                       ************************************************************

alpha1_core_sheep      RIRKLADQCTGLQGFLVFHSFGGGTGSGFTSLLMERLSVDYGKKSKLEFSIYPAPQVSTA
alpha1_6CVN            RIRKLADQCTGLQGFLVFHSFGGGTGSGFTSLLMERLSVDYGKKSKLEFSIYPAPQVSTA
                       ************************************************************

alpha1_core_sheep      VVEPYNSILTTHTTLEHSDCAFMVDNEAIYDICRRNLDIERPTYTNLNRLIGQIVSSITA
alpha1_6CVN            VVEPYNSILTTHTTLEHSDCAFMVDNEAIYDICRRNLDIERPTYTNLNRLISQIVSSITA
                       ***************************************************.********

alpha1_core_sheep      SLRFDGALNVDLTEFQTNLVPYPRIHFPLATYAPVISAEKAYHEQLSVAEITNACFEPAN
alpha1_6CVN            SLRFDGALNVDLTEFQTNLVPYPRIHFPLATYAPVISAEKAYHEQLSVAEITNACFEPAN
                       ************************************************************

alpha1_core_sheep      QMVKCDPRHGKYMACCLLYRGDVVPKDVNAAIATIKTKRTIQFVDWCPTGFKVGINYQPP
alpha1_6CVN            QMVKCDPRHGKYMACCLLYRGDVVPKDVNAAIATIKTKRSIQFVDWCPTGFKVGINYQPP
                       ***************************************:********************

alpha1_core_sheep      TVVPGGDLAKVQRAVCMLSNTTAIAEAWARLDHKFDLMYAKRAFVHWYVGEGMEEGEFSE
alpha1_6CVN            TVVPGGDLAKVQRAVCMLSNTTAIAEAWARLDHKFDLMYAKRAFVHWYVGEGMEEGEFSE
                       ************************************************************

alpha1_core_sheep      AREDMAALEKDYEEVGVD
alpha1_6CVN            AREDMAALEKDYEEVGVD
                       ******************
```

```
beta1_core_sheep      MREIVHIQAGQCGNQIGAKFWEVISDEHGIDPTGTYHGDSDLQLDRISVYYNEATGGKYV
beta1_6CVN            MREIVHIQAGQCGNQIGAKFWEVISDEHGIDPTGSYHGDSDLQLERINVYYNEAAGNKYV
                      **********************************:*********:**.******:*.***

beta1_core_sheep      PRAILVDLEPGTMDSVRSGPFGQIFRPDNFVFGQSGAGNNWAKGHYTEGAELVDSVLDVV
beta1_6CVN            PRAILVDLEPGTMDSVRSGPFGQIFRPDNFVFGQSGAGNNWAKGHYTEGAELVDSVLDVV
                      ************************************************************

beta1_core_sheep      RKEAESCDCLQGFQLTHSLGGGTGSGMTLLISKIREEYPDRIMNTFSVVPSPKVSDTVV
beta1_6CVN            RKESESCDCLQGFQLTHSLGGGTGSGMTLLISKIREEYPDRIMNTFSVVPSPKVSDTVV
                      ***:*******************************************************

beta1_core_sheep      EPYNATLSVHQLVENTDETYCIDNEALYDICFRTLKLTTPTYGDLNHLVSATMSGVTTCL
beta1_6CVN            EPYNATLSVHQLVENTDETYCIDNEALYDICFRTLKLTTPTYGDLNHLVSATMSGVTTCL
                      ************************************************************

beta1_core_sheep      RFPGQLNADLRKLAVNMVPFPRLHFFMPGFAPLTSRGSQQYRALTVPELTQQVFDAKNMM
beta1_6CVN            RFPGQLNADLRKLAVNMVPFPRLHFFMPGFAPLTSRGSQQYRALTVPELTQQMFDAKNMM
                      ****************************************************:*******

beta1_core_sheep      AACDPRHGRYLTVAAVFRGRMSMKEVDEQMLNVQNKNSSYFVEWIPNNVKTAVCDIPPRG
beta1_6CVN            AACDPRHGRYLTVAAVFRGRMSMKEVDEQMLNVQNKNSSYFVEWIPNNVKTAVCDIPPRG
                      ************************************************************

beta1_core_sheep      LKMAVTFIGNSTAIQELFKRISEQFTAMFRRKAFLHWYTGEGMDEMEFTEAESNMNDLVS
beta1_6CVN            LKMSATFIGNSTAIQELFKRISEQFTAMFRRKAFLHWYTGEGMDEMEFTEAESNMNDLVS
                      ***:.*******************************************************

beta1_core_sheep      EYQQYQ
beta1_6CVN            EYQQYQ
                      ******


beta3_core_human      MREIVHIQAGQCGNQIGAKFWEVISDEHGIDPSGNYVGDSDLQLERISVYYNEASSHKYV
beta1_6CVN            MREIVHIQAGQCGNQIGAKFWEVISDEHGIDPTGSYHGDSDLQLERINVYYNEAAGNKYV
                      ********************************:*.* **********.******:.:***

beta3_core_human      PRAILVDLEPGTMDSVRSGAFGHLFRPDNFIFGQSGAGNNWAKGHYTEGAELVDSVLDVV
beta1_6CVN            PRAILVDLEPGTMDSVRSGPFGQIFRPDNFVFGQSGAGNNWAKGHYTEGAELVDSVLDVV
                      ******************* **::******:*****************************

beta3_core_human      RKECENCDCLQGFQLTHSLGGGTGSGMTLLISKVREEYPDRIMNTFSVVPSPKVSDTVV
beta1_6CVN            RKESESCDCLQGFQLTHSLGGGTGSGMTLLISKIREEYPDRIMNTFSVVPSPKVSDTVV
                      ***.*.***************************:*************************

beta3_core_human      EPYNATLSIHQLVENTDETYCIDNEALYDICFRTLKLATPTYGDLNHLVSATMSGVTTSL
beta1_6CVN            EPYNATLSVHQLVENTDETYCIDNEALYDICFRTLKLTTPTYGDLNHLVSATMSGVTTCL
                      ********:****************************:********************.*

beta3_core_human      RFPGQLNADLRKLAVNMVPFPRLHFFMPGFAPLTARGSQQYRALTVPELTQQMFDAKNMM
beta1_6CVN            RFPGQLNADLRKLAVNMVPFPRLHFFMPGFAPLTSRGSQQYRALTVPELTQQMFDAKNMM
                      **********************************:*************************

beta3_core_human      AACDPRHGRYLTVATVFRGRMSMKEVDEQMLAIQSKNSSYFVEWIPNNVKVAVCDIPPRG
beta1_6CVN            AACDPRHGRYLTVAAVFRGRMSMKEVDEQMLNVQNKNSSYFVEWIPNNVKTAVCDIPPRG
                      **************:**************** :*.***************.*********

beta3_core_human      LKMSSTFIGNSTAIQELFKRISEQFTAMFRRKAFLHWYTGEGMDEMEFTEAESNMNDLVS
beta1_6CVN            LKMSATFIGNSTAIQELFKRISEQFTAMFRRKAFLHWYTGEGMDEMEFTEAESNMNDLVS
                      ****:*******************************************************

beta3_core_human      EYQQYQ
beta1_6CVN            EYQQYQ
                      ******
```

S3

**Figure SI-1:** DSSP graphs for tau-R2 along all the MD trajectories.
(a-c) tau-R2 in solution, (d-f) tau-R2 bound on tubulin without CTTs.

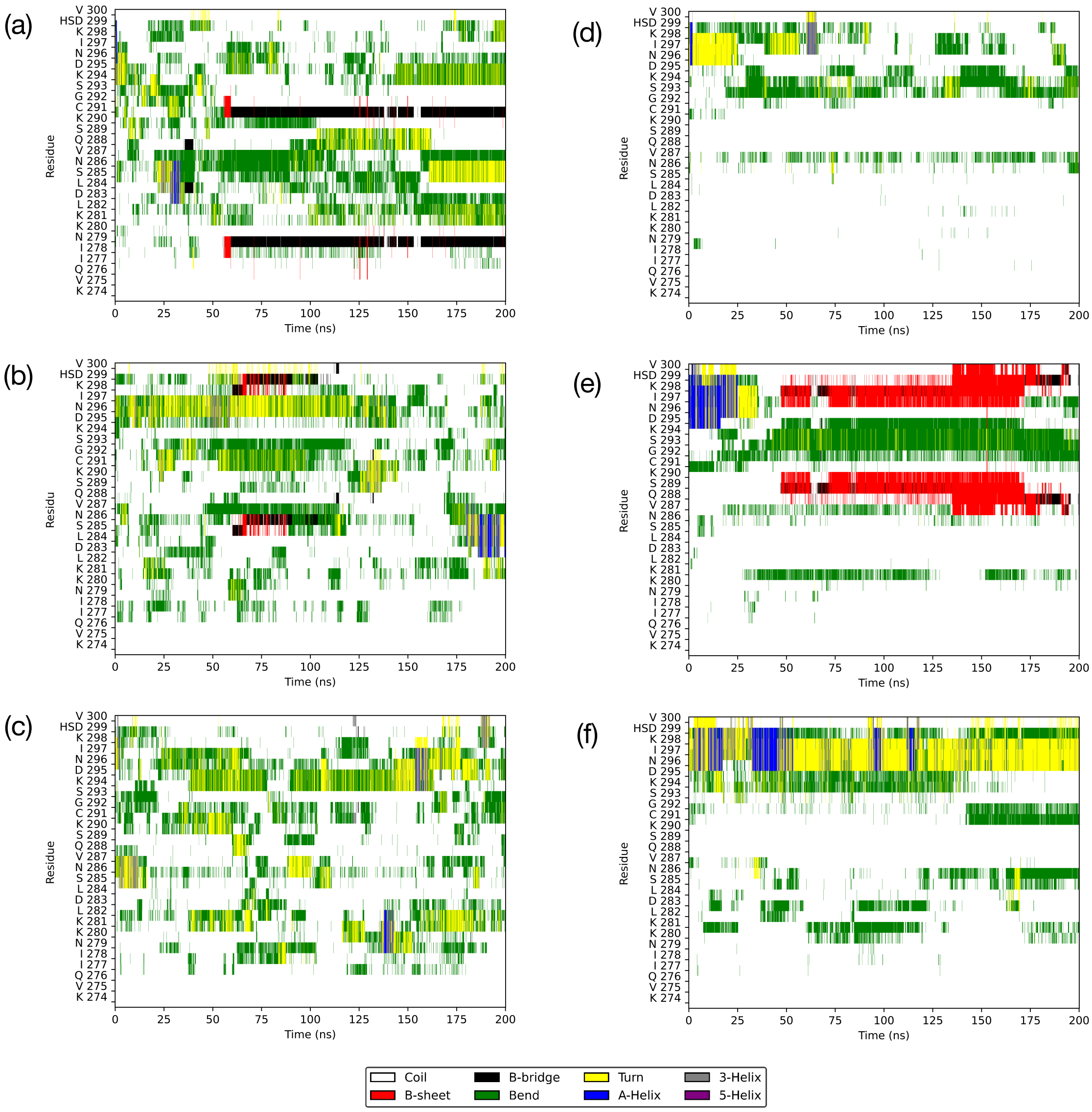

**Figure SI-1(continued):** DSSP graphs for tau-R2 along all the MD trajectories.
(g-i) tau-R2 bound to the βI/αI/βI isotype, (j-l) tau-R2 bound to the βIII/αI/βIII isotype.

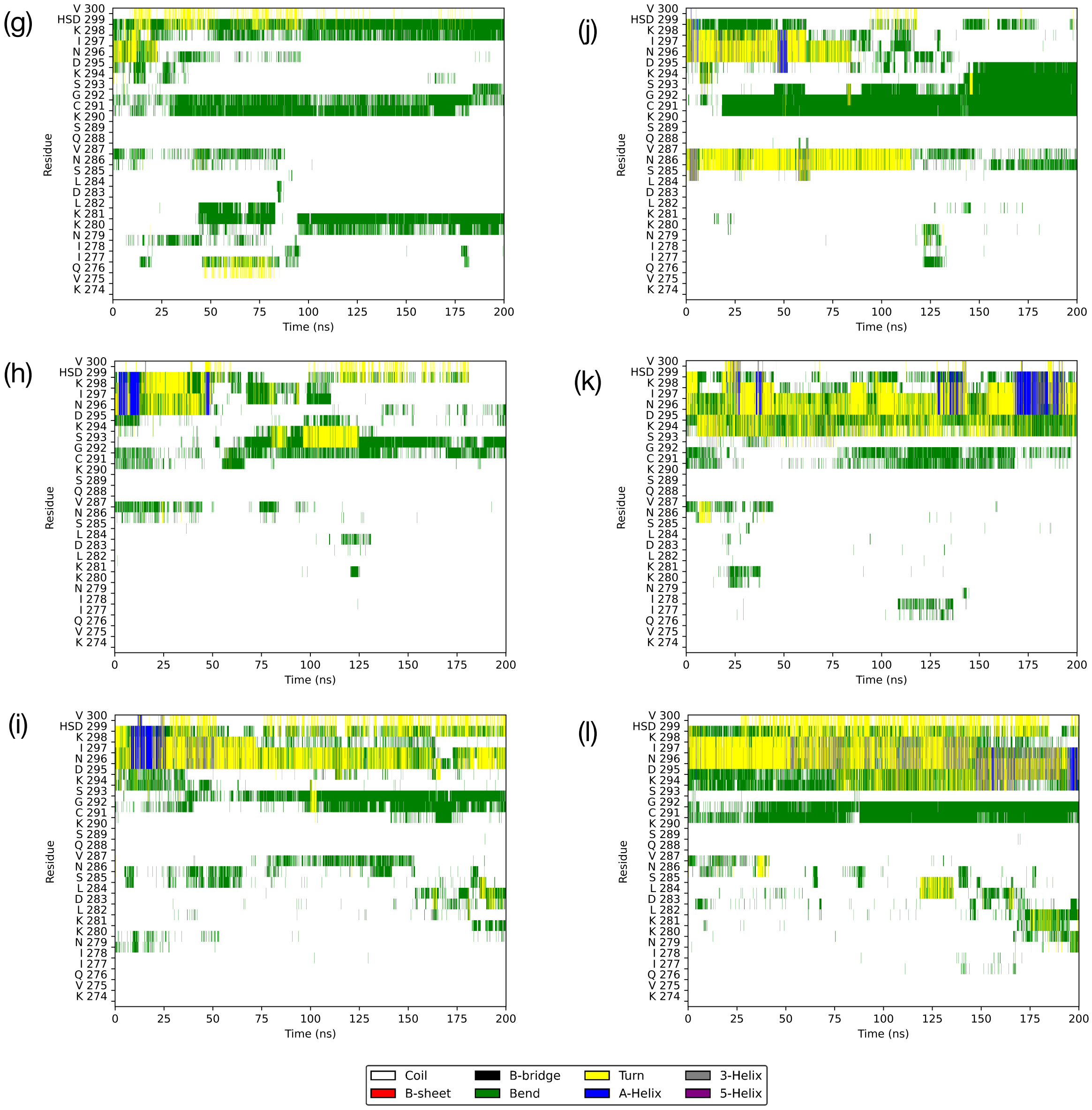

**Figure SI-2I:** DSSP graphs for the tubulin CTTs along the MD trajectories, βI/αI/βI isotype without tau-R2.
Model 17: (a) first β-subunit, (c) α-subunit, (d) second β-subunit.
Model 22: (d) first β-subunit, (e) α-subunit, (f) second β-subunit.
Model 84: (g) first β-subunit, (h) α-subunit, (i) second β-subunit.

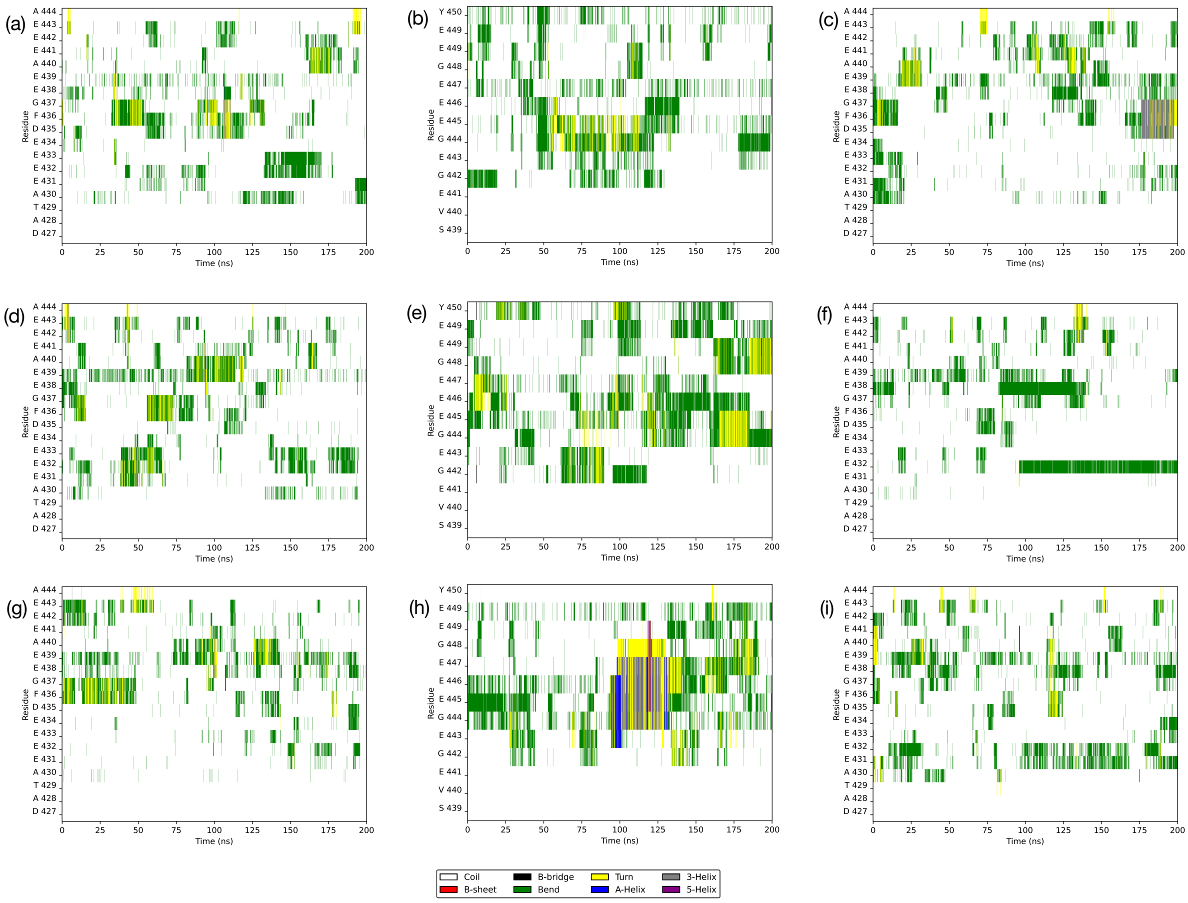

**Figure SI-2II:** DSSP graphs for the tubulin CTTs along the MD trajectories, βI/αI/βI isotype with tau-R2.
Model 17: (a) first β-subunit, (c) α-subunit, (d) second β-subunit.
Model 22: (d) first β-subunit, (e) α-subunit, (f) second β-subunit.
Model 84: (g) first β-subunit, (h) α-subunit, (i) second β-subunit.

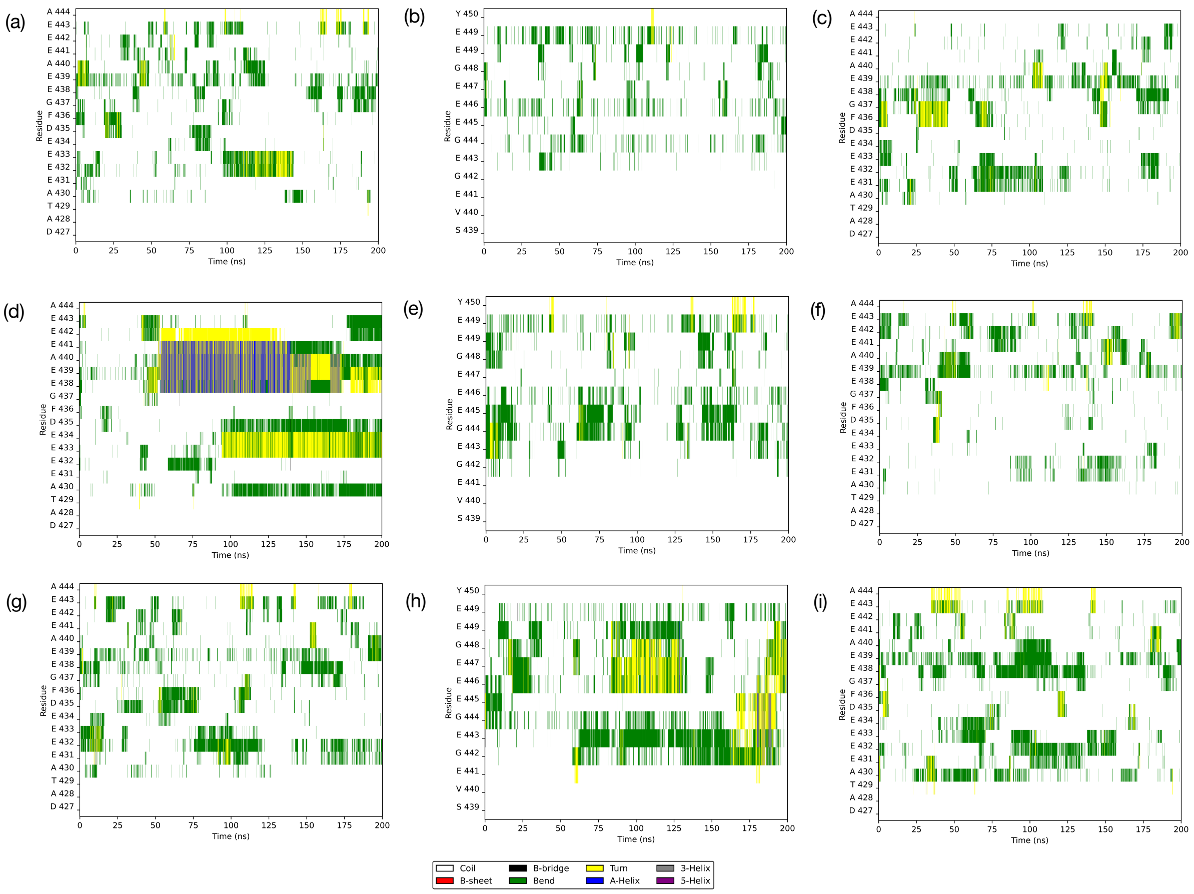

**Figure SI-2III:** DSSP graphs for the tubulin CTTs along the MD trajectories, βIII/αI/βIII isotype without tau-R2.
Model 16: (a) first β-subunit, (c) α-subunit, (d) second β-subunit.
Model 38: (d) first β-subunit, (e) α-subunit, (f) second β-subunit.
Model 65: (g) first β-subunit, (h) α-subunit, (i) second β-subunit.

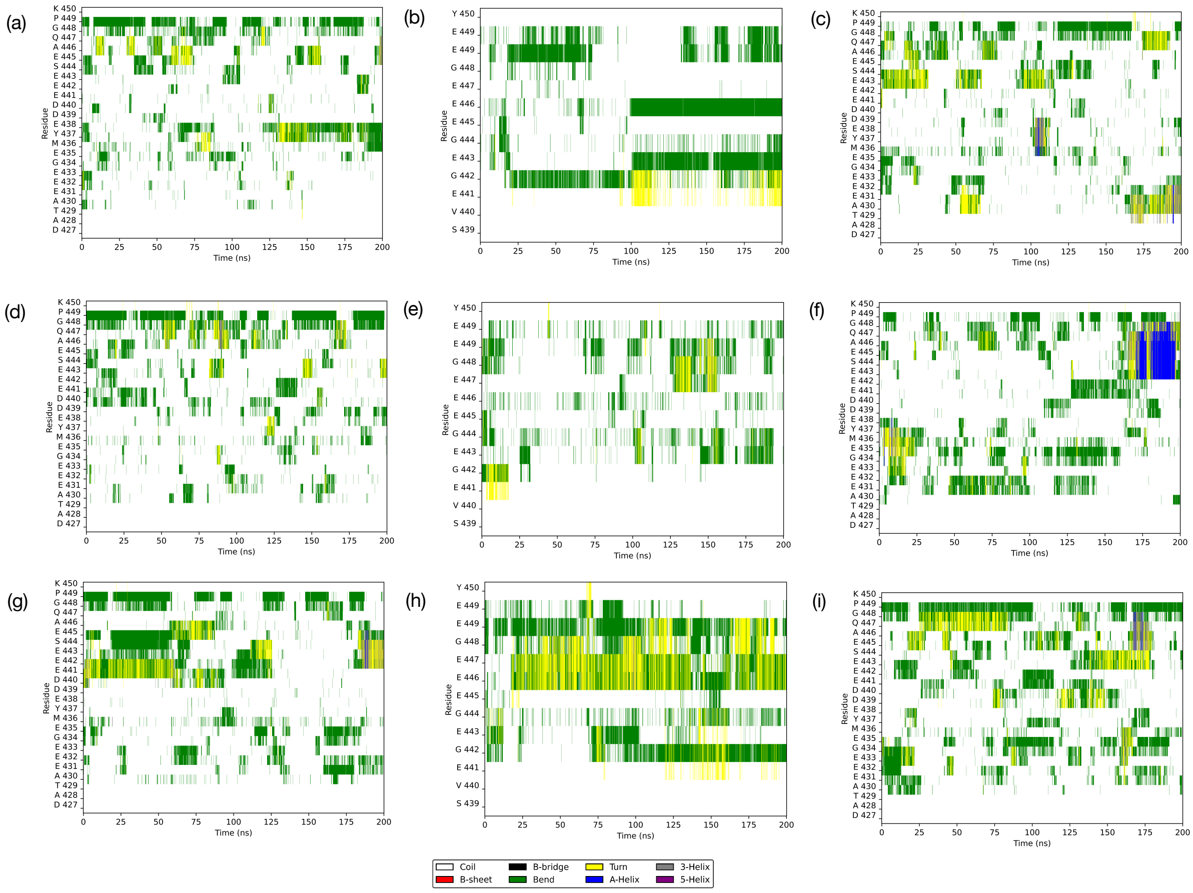

**Figure SI-2IV:** DSSP graphs for the tubulin CTTs along the MD trajectories, βIII/αI/βIII isotype with tau-R2.
Model 16: (a) first β-subunit, (c) α-subunit, (d) second β-subunit.
Model 38: (d) first β-subunit, (e) α-subunit, (f) second β-subunit.
Model 65: (g) first β-subunit, (h) α-subunit, (i) second β-subunit.

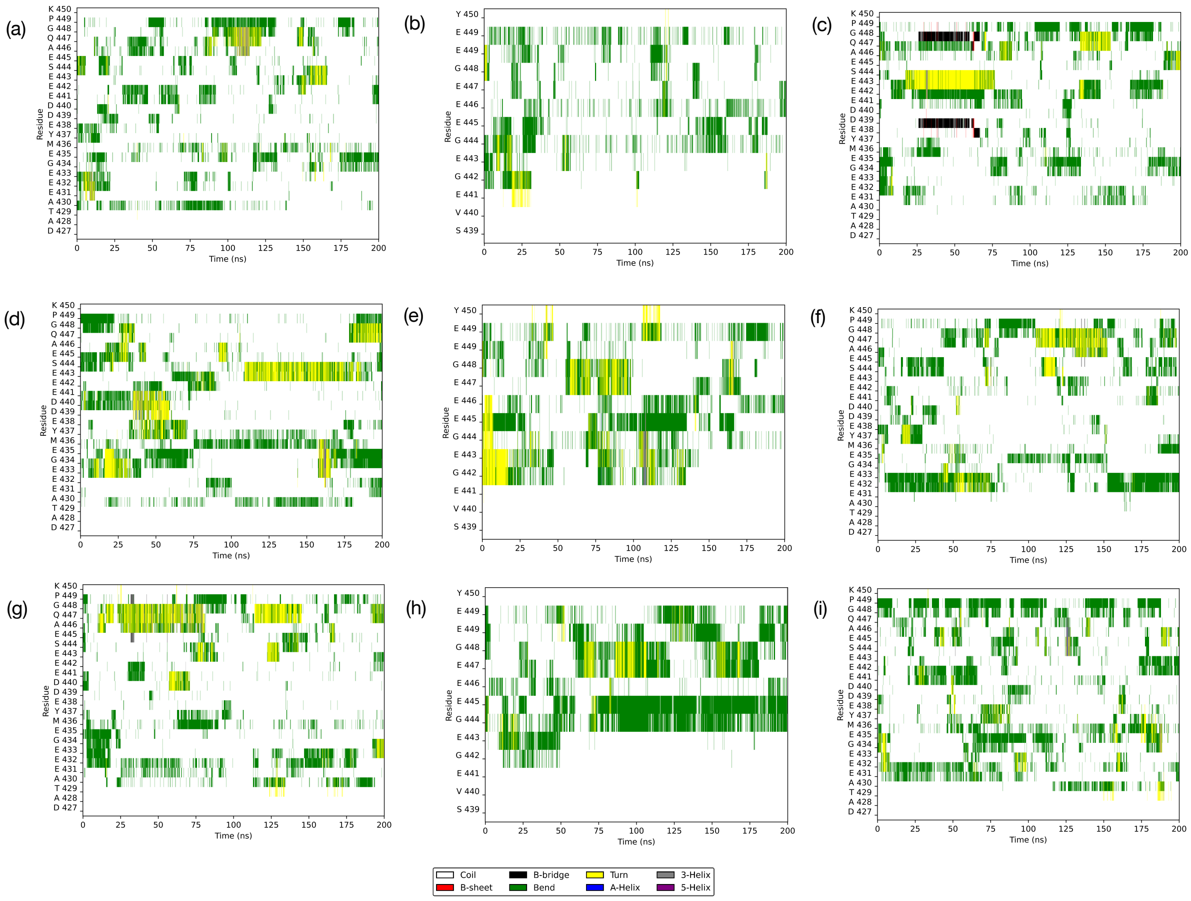

**Figure SI-3:** Backbone RMSDs (smoothed using a 50 frames, 5 ns, moving window, while conserving the detailed data as transparent background) as a function of time for the three MD simulations of tau-R2 in solution.

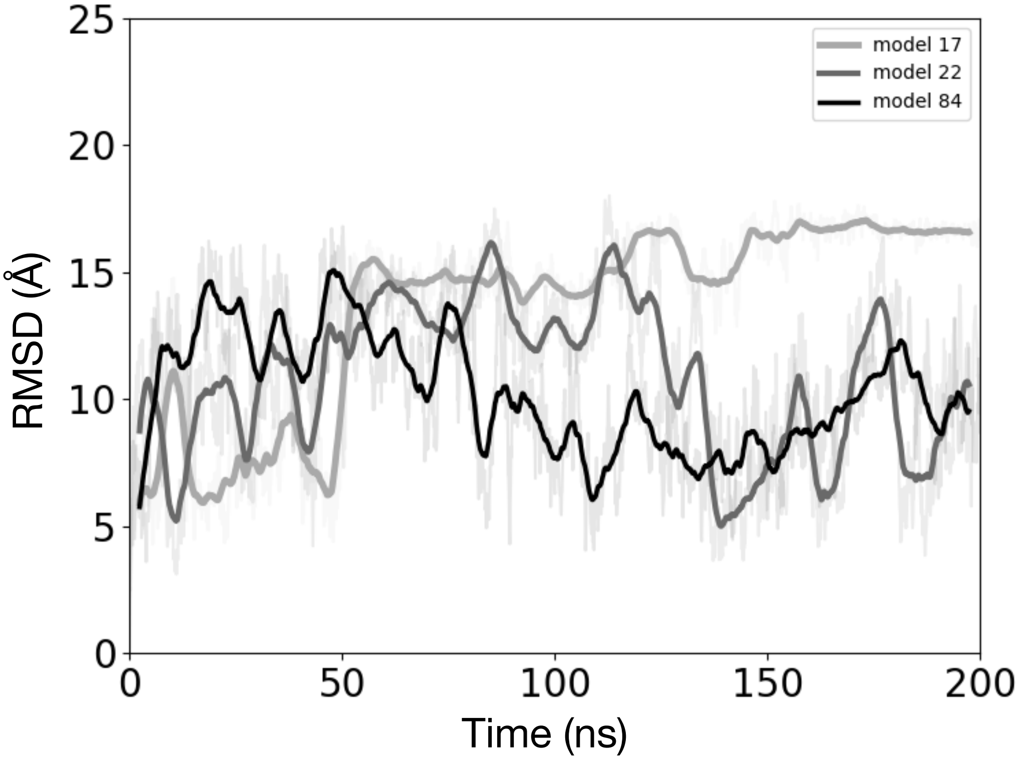

**Figure 4:** Backbone RMSDs (smoothed using a 50 frames, 5 ns, moving window, while conserving the detailed data as transparent background) as a function of time for the tau-R2/tubulin complexes. Black and grey lines for the tubulin without CTTs, blue lines for the βI/αI/βI isotype and red-orange lines for the βIII/αI/βIII isotype (this color code is also used for Figures SI-5-7 and 10). (a)-(c) Whole system, (d)-(f) tubulin core only, (g)-(i) tau-R2 only, (j)-(k) CTTs only.

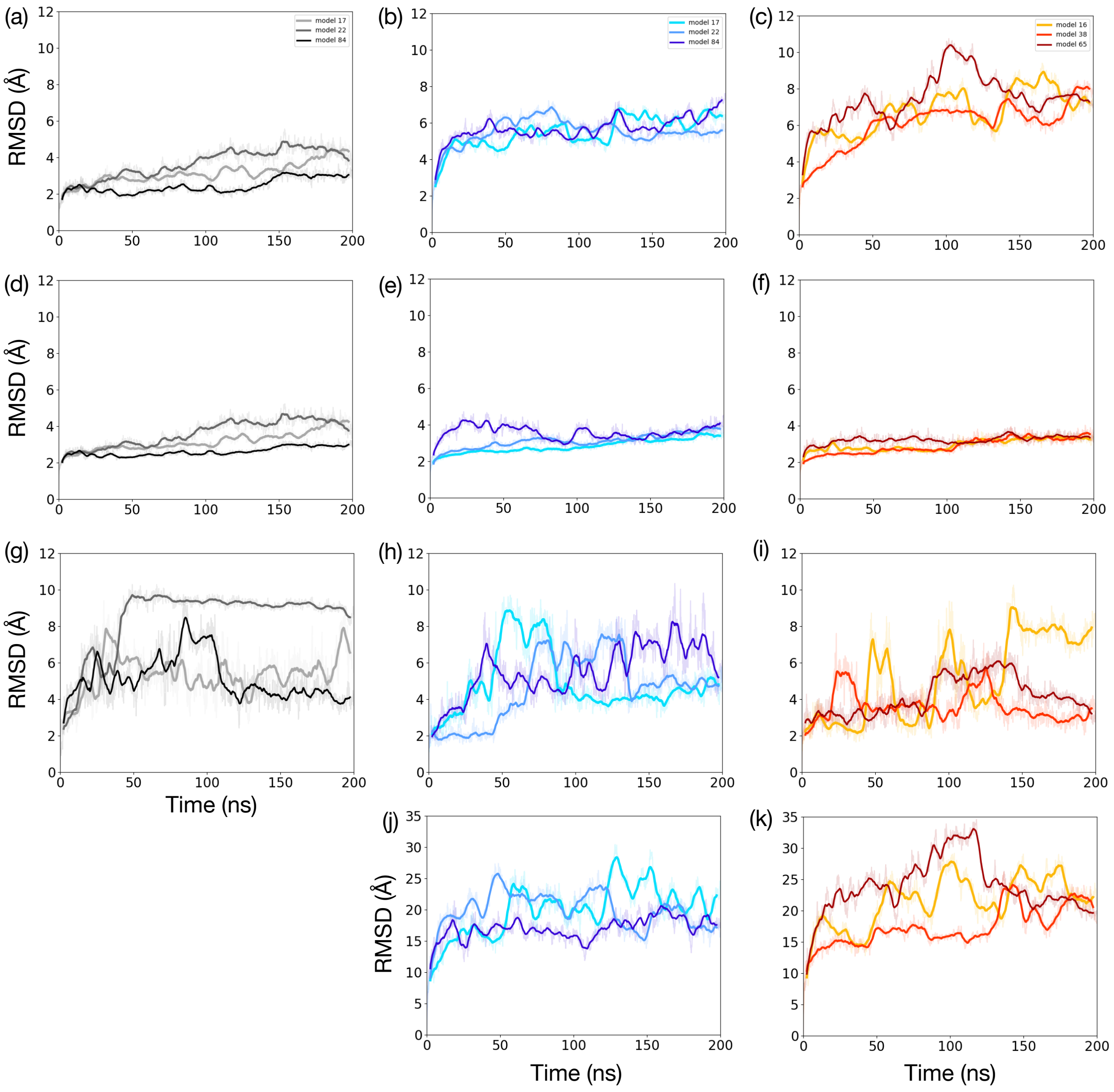

**Figure SI-5:** Fraction of native contacts between tau-R2 and the tubulin heterotrimer as a function of time (a) Tubulin without CTTs (b) βI/αI/βI isotype (c) βIII/αI/βIII isotype.

Total number of contacts between tau-R2 and the tubulin heterotrimer as a function of time (d) Tubulin without CTTs (e) βI/αI/βI isotype (f) βIII/αI/βIII isotype.

All curves were smoothed using a 50 frames, 5 ns, moving window, while conserving the detailed data as transparent background.

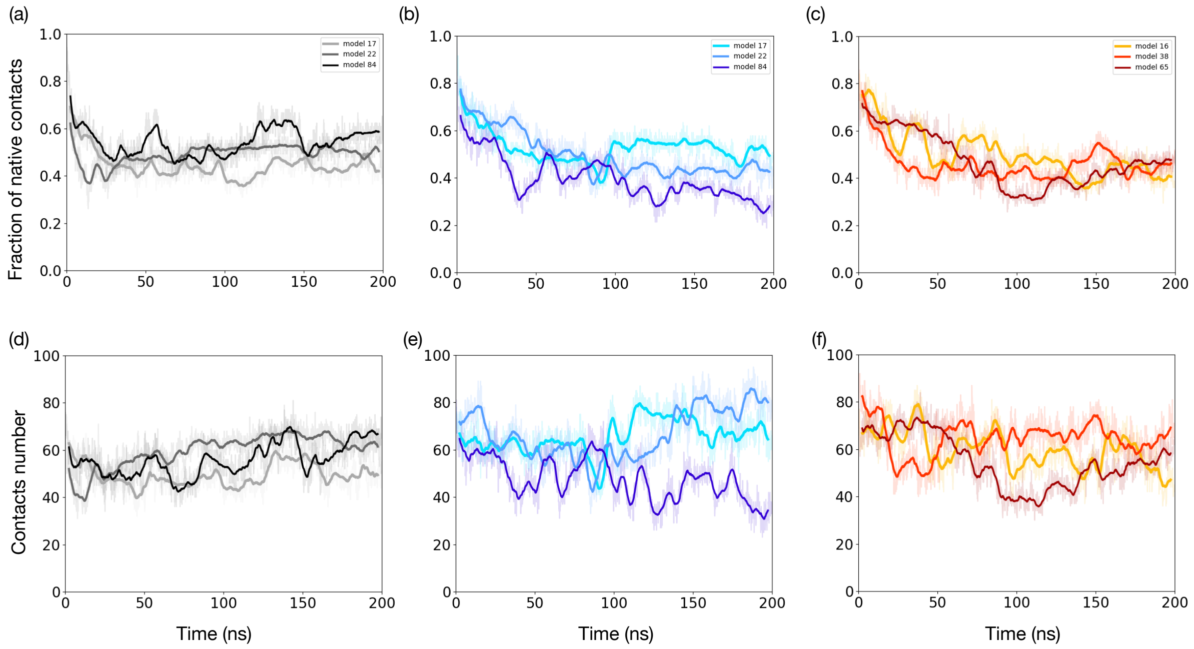

**Figure SI-6:** RMSF of tau-R2, for the nine MD simulations: (top) Tubulin without CTTs (center) βI/αI/βI isotype (bottom) βIII/αI/βIII isotype. The three serine residues that are phosphorylated in tauopathies are highlighted in green along the sequence.

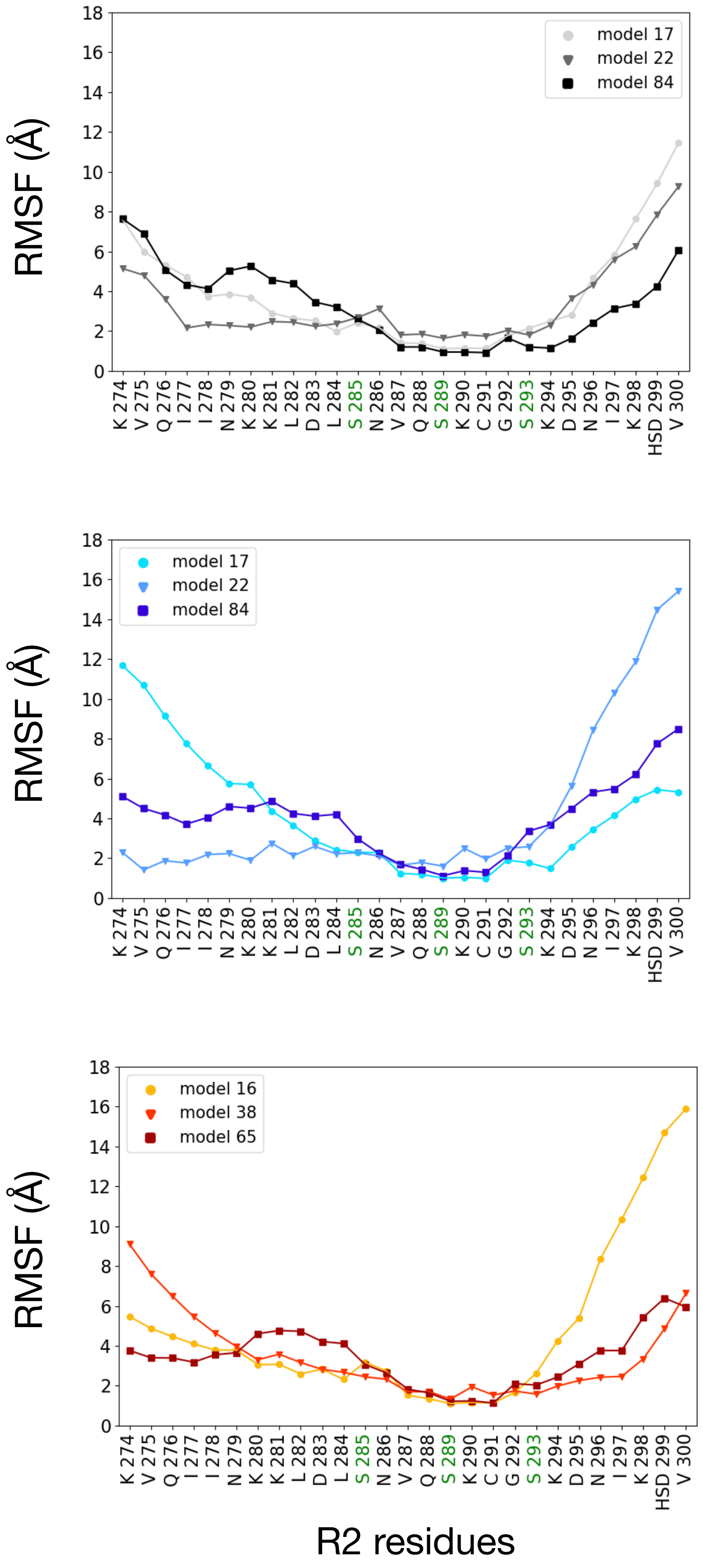

**Figure SI-7:** Distribution of the number of hydrogen bonds formed by the three serine residues of tau-R2 during the MD trajectories for the various systems modeled (tau-R2 in solution, in complex with tubulins without CTTs, in complex with the βI/αI/βI isotype, in complex with the βIII/αI/βIII isotype).

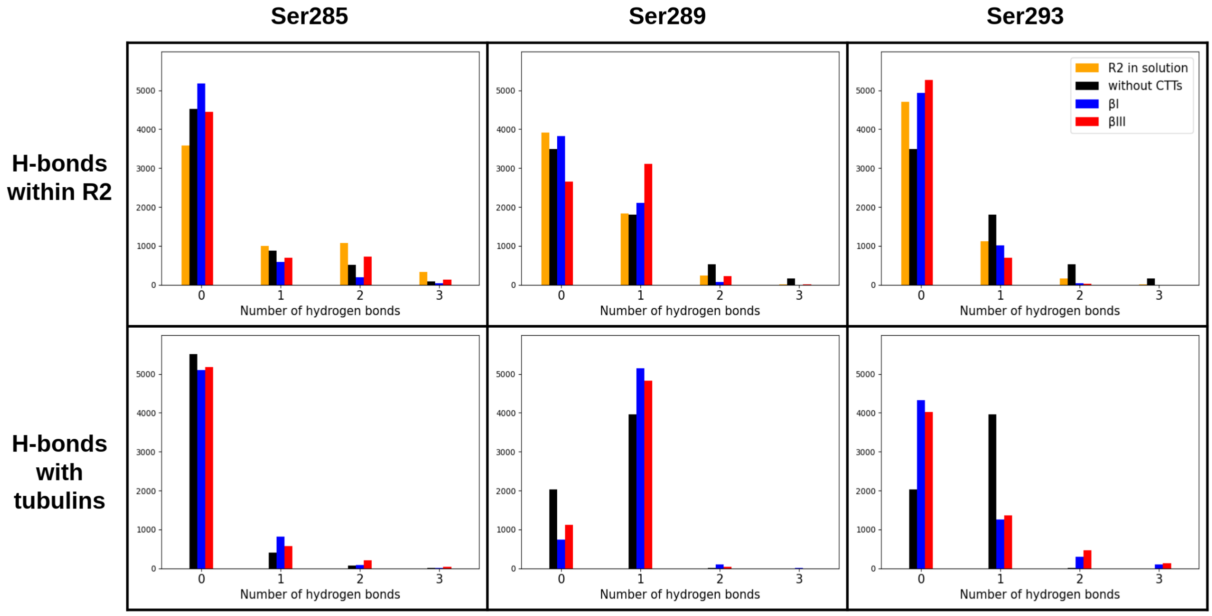

**Figure SI-8:** Mobility pattern of the CTTs center of mass during the MD trajectories (three replicas of 200 ns for each system) of the tubulin heterotrimer without (left panels) and with bound tau-R2 (right panels). (top) βI/αI/βI isotype, (bottom) βIII/αI/βIII isotype. The tubulin heretotrimer is shown as a transparent cartoon in the background and the initial position of tau-R2 is shown in black as a visual guide. The inserts above the mobility distribution plots show the structure of a CTT (in orange) corresponding to the circled area.

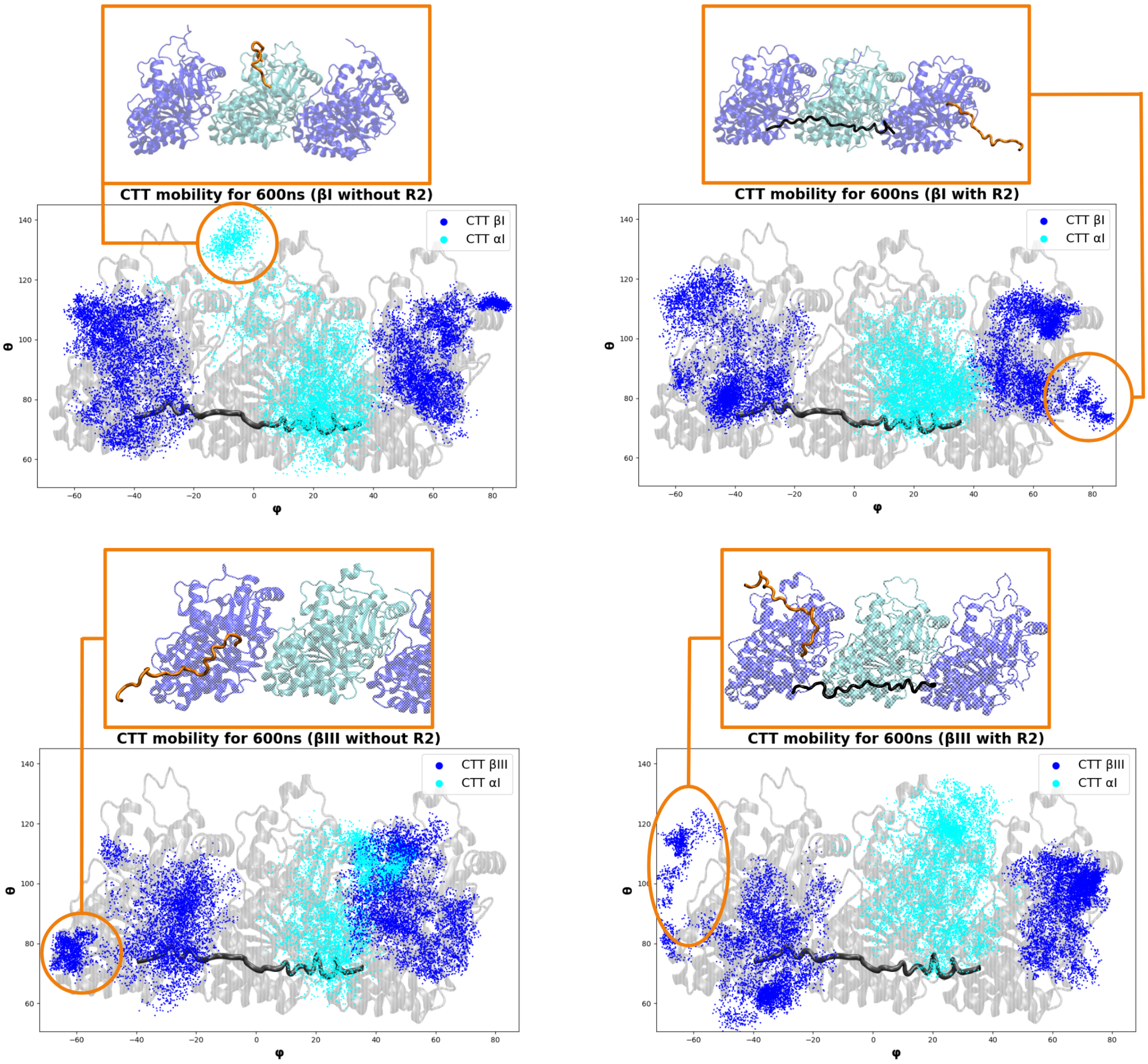

**Figure SI-9:** Probability density for the number of contacts formed by each tubulin-CTT

Tubulin heterotrimer without tau-R2, contacts formed between the CTT and the tubulin core:

- βI/αI/βI isotype, (a) First β-subunit, (b) α-subunit (c) second β-subunit
- βIII/αI/βIII isotype, (d) First β-subunit, (e) α-subunit (f) second β-subunit

Tubulin heterotrimer with tau-R2, contacts formed between the CTT and the tubulin core or between the CTT and tau-R2:

- βI/αI/βI isotype, (g) First β-subunit, (h) α-subunit (i) second β-subunit
- βIII/αI/βIII isotype, (j) First β-subunit, (k) α-subunit (l) second β-subunit

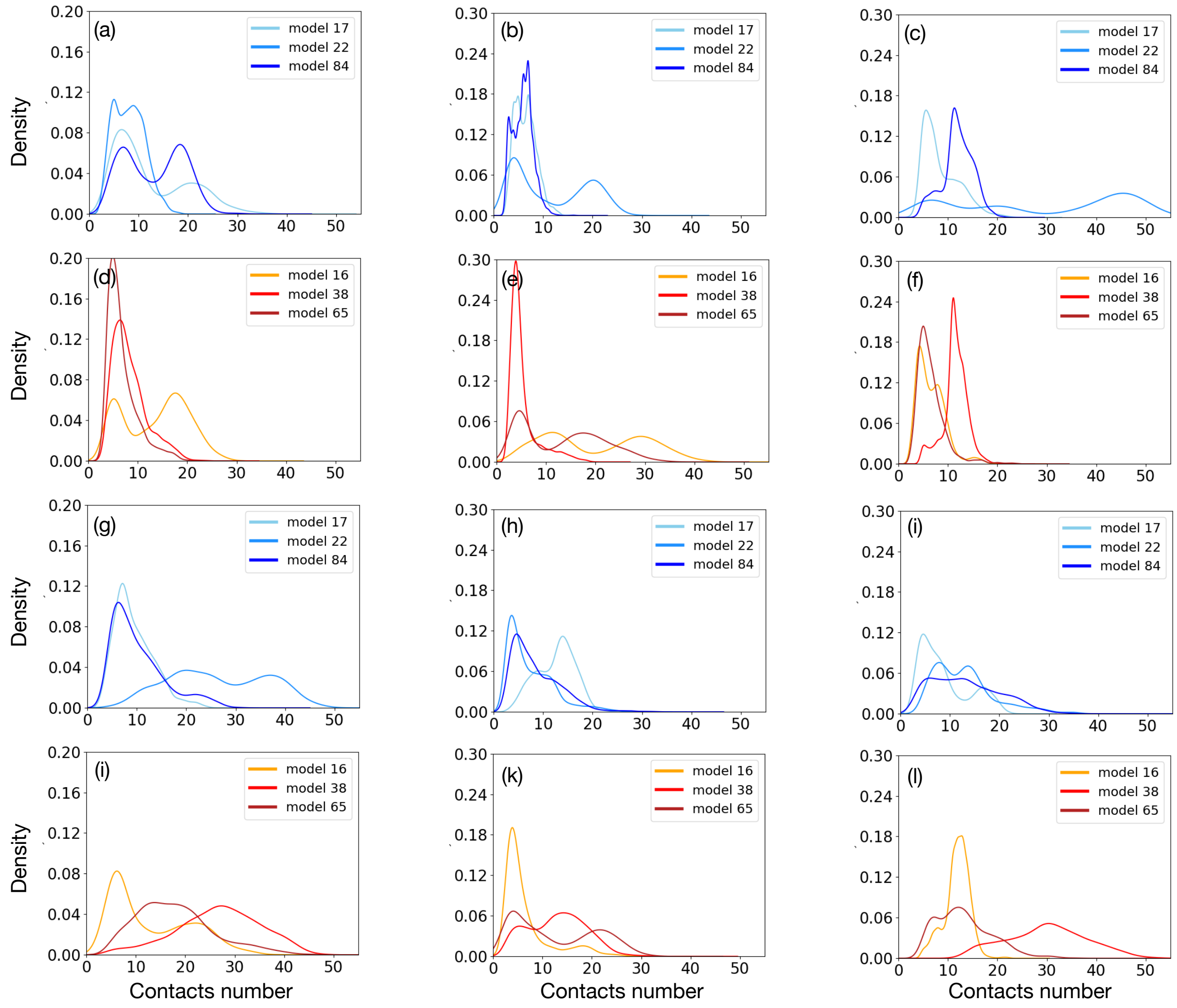

**Figure SI-10:** Number of contacts formed by each CTT in the βI/αI/βI isotype as a function of time :

- Simulations without tau-R2, contacts formed between the CTT and the tubulin core (a) First β-subunit, (b) α-subunit (c) second β-subunit.
- Simulations with tau-R2, contacts formed between the CTT and the tubulin core or between the CTT and tau-R2 (d) First β-subunit, (e) α-subunit (f) second β-subunit.

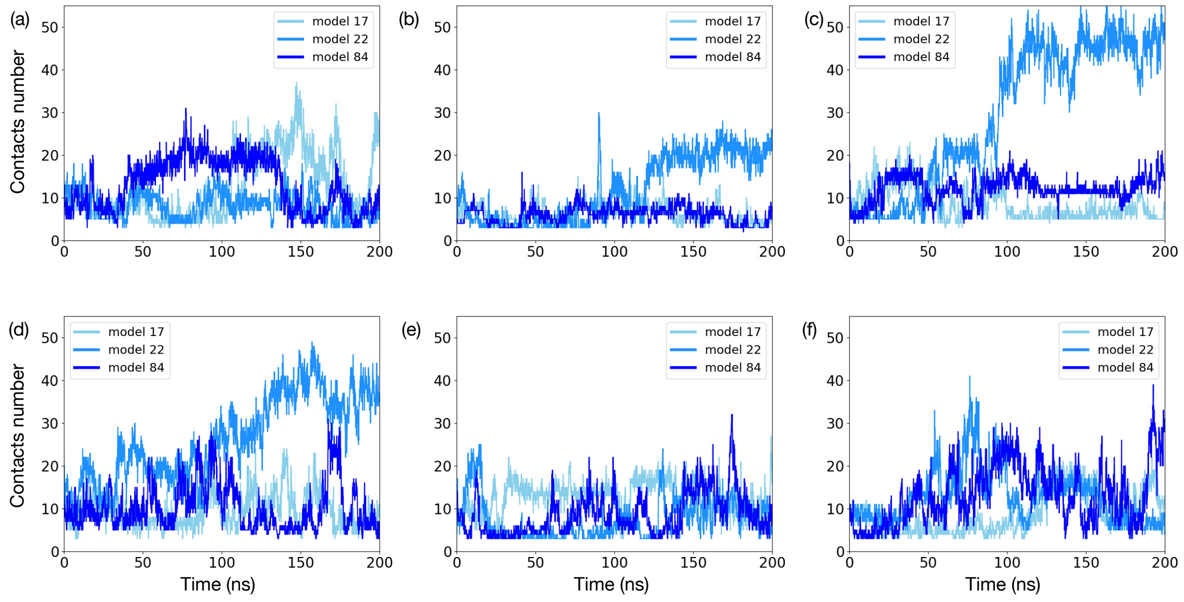

**Figure SI-11:** Number of contacts formed by each CTT in the βIII/αI/βIII isotype as a function of time :

- Simulations without tau-R2, contacts formed between the CTT and the tubulin core (a) First β-subunit, (b) α-subunit (c) second β-subunit.

- Simulations with tau-R2, contacts formed between the CTT and the tubulin core or between the CTT and tau-R2 (d) First β-subunit, (e) α-subunit (f) second β-subunit.

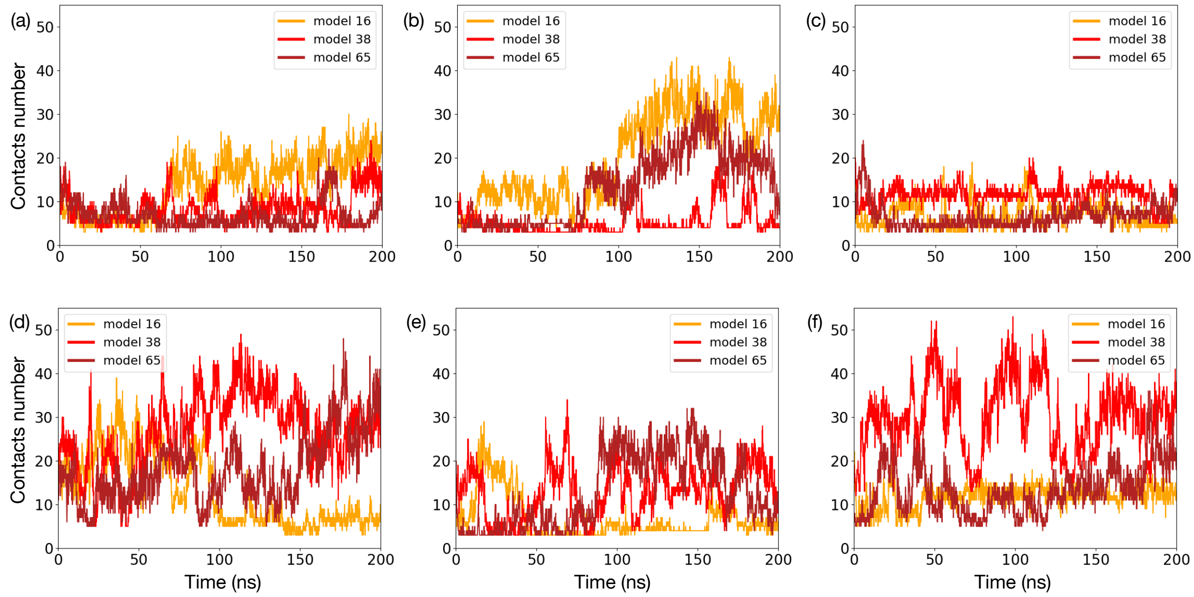
